# A sequence-encoded promoter proximal super pause stabilizes an offline RNA polymerase II state

**DOI:** 10.64898/2026.02.18.706689

**Authors:** Roberto J. Vazquez Nunez, Serena Kesha, Seychelle M. Vos

**Affiliations:** Department of Biology, Massachusetts Institute of Technology; Cambridge, 02139, USA; Howard Hughes Medical Institute

## Abstract

Promoter proximal pausing by RNA polymerase II is critical for regulating gene expression in multicellular eukaryotes. How nucleic acid sequence and protein factors contribute to pausing remains incompletely understood. We developed Gene-specific Analysis of Transcriptional Output (GATO)-seq, which for the first time enables massively parallel, temporally resolved, reconstituted transcription in an assay that uses direct RNA sequencing to map 3′ends of nascent transcripts from a library of human genes. GATO-seq identified a “super pause” sequence that potently induces RNA polymerase II pausing and is not relieved by rescue factor Transcription Factor (TF) IIS. Cryogenic-electron microscopy (cryo-EM) structures of RNA polymerase II on the super pause sequence reveal a previously unobserved, reversible single-nucleotide backtracked state (“sidetracked”), stabilized by a threonine-lined pocket that limits further backtracking. We introduce a powerful in vitro technique that can be employed to study transcription regulation and through its use show that nucleic acid sequence encodes pausing propensity and traps sequence specific offline states, linking sequence to pausing control.

## Introduction

Transcription is tightly regulated to ensure appropriate gene expression. RNA polymerase II transcribes most eukaryotic protein-coding genes and its activity is regulated by transcription factors, the underlying DNA sequence, chromatin, and co-transcriptional events such as splicing ^1–3^. RNA polymerase II is prone to pause in the promoter-proximal region of genes^4,5^. Promoter-proximal pausing occurs on nearly all genes and serves multiple regulatory functions, including enabling rapid and coordinated transcriptional responses, maintaining open chromatin at promoter regions, and acting as a quality-control checkpoint to ensure proper assembly of competent elongation complexes^6–13^.

Regulation of promoter proximal pausing in metazoans involves several protein complexes that associate with RNA polymerase II. DRB-sensitivity inducing factor (DSIF) and Negative Elongation Factor (NELF) bind RNA polymerase II to form stably paused elongation complexes^14–19^. Promoter-proximal paused RNA polymerases can either be released into productive transcription elongation or prematurely terminated. Release into productive elongation is mediated by the kinase positive transcription elongation factor b (P-TEFb), which phosphorylates the RNA polymerase II subunit RPB1 C-terminal domain (CTD) and CTD linker, as well as DSIF-NELF^20–27^. These modifications help displace NELF and promote recruitment of elongation factors SPT6 and PAF1c to form the activated elongation complex (EC*) with DSIF^23,28^. Premature termination can involve the Integrator complex, which binds the RNA polymerase II-DSIF-NELF complex^29–35^. Integrator is thought to evict RNA polymerase II from DNA through its combined phosphatase and RNA endonuclease activities. Additional factors, including Facilitates Chromatin Transcription (FACT), ARMC5, Transcription Factor (TF) IID, and the +1 nucleosome are implicated in regulating promoter-proximal pausing^13,36–41^.

In addition to protein regulators, the underlying DNA sequence at promoter-proximal regions plays an important role in pausing. Cell based assays including Native Elongating Transcript sequencing (NET-seq) and Precision-Run-On sequencing (PRO-seq) map RNA polymerase position at single nucleotide resolution by sequencing RNAs associated with RNA polymerase or through the incorporation of biotinylated nucleotide triphosphates (NTPs)^42–44^. These experiments have reported RNA polymerase II enrichment on specific sequences (**Fig. S1A**)^42,44–51^. Notably, some of these sequence features overlap with those of the bacterial elemental pause sequence. Mammalian RNA polymerase II and even the evolutionarily distal human mitochondrial RNA polymerase have been shown to pause at the bacterial elemental pause sequence, suggesting that fundamental DNA-encoded mechanisms of pausing may be conserved across domains of life^45,52–56^.

Most cell-based assays are performed under steady-state conditions and provide limited temporal resolution, making it difficult to assess pause duration, frequency, and the precise contributions of individual protein factors to pausing, pause release, and premature termination. Single-molecule and bulk biochemical experiments facilitate tight temporal control and have shown that specific sequences can modulate pause duration^52,57–62^, but are usually obtained from experiments performed at nucleotide concentrations far below those found in cells and have been restricted to a handful of model sequences^55,58,59^, limiting our understanding of how nucleic acid sequence may impact pausing in a biologically relevant context. Thus, improved reconstituted approaches are urgently needed to better recapitulate the conditions found in cells and assess transcriptional pausing activity on a wide range of DNA sequences.

Here we present GATO-seq (Gene-specific Analysis of Transcriptional Output), a high-throughput transcription assay developed to overcome these limitations, and study transcription in a sequence, time, and factor dependent manner at single nucleotide and single molecule resolution using highly purified proteins. To establish GATO-seq, we examined how DNA sequence affects RNA polymerase II elongation using a library of human promoter-proximal sequences. We confirm that RNA polymerase II pause duration is distinct on different DNA sequences and that nucleotide concentration and protein factors regulate translocation speed and release from paused states. Pause sites that are refractory to rescue factor TFIIS share a conserved consensus sequence and exhibit exceptionally long dwell times, which we term “super pauses”. Cryo-EM structures of RNA polymerase II on the consensus super pause sequence reveals, to our knowledge, a previously unobserved, reversible, one-nucleotide backtracked register, termed sidetracking, stabilized by a threonine-lined pocket. Together, our results demonstrate that GATO-seq can disentangle DNA-encoded and factor-dependent contributions to transcriptional pausing and uncovers a new offline state that may function as a regulatory element in promoter-proximal regions of genes.

## Results

### GATO-seq reports on elongation velocity

Our goal was to establish a cell-free genomics approach to generally study transcription using highly purified proteins and nucleic acids. As a proof of principle, we investigated how sequence influences RNA polymerase II pausing behavior. This was implemented on a DNA library containing 1,000 human genes with well-focused transcription start sites (TSS) that are associated with various degrees of promoter-proximal pausing in HEK293T cells (**Fig. S1B, Supplemental Table 1**). We used the first 262 base pairs (bp) of each gene because these sequences are most associated with RNA polymerase II pausing behaviors in cells. To eliminate variability arising from promoter usage and to allow for kinetic analysis of transcription elongation, the first 262 base pairs of each gene were fused downstream of an Adenovirus Major Late (AdML) promoter, a 9 bp G-less cassette and a 10 bp adapter sequence. This results in a maximum transcript length of 300 bp (**Fig. 1A**). Promoter sequence influences early transcription elongation behavior and thus our assay reports on general elongation behaviors independent of promoter context^37,50,63^. Pre-initiation complexes consisting of highly purified porcine RNA polymerase II and human Transcription Factor (TF) IIA, TFIIB, TFIIE, TFIIF, TFIIH, Cyclin Dependent Kinase activating kinase (CAK) and TATA-binding protein (TBP) were assembled on the library (**Fig. 1A, B, Fig. S1C**). Transcription was initiated and permitted to proceed through the G-less cassette by adding ATP, CTP, and UTP. To synthesize RNA past the G-less cassette and prevent re-initiation, GTP and triptolide, an inhibitor of the TFIIH helicase XPB^64^, were added to the reaction, respectively. Reactions were quenched at various time points after GTP addition, and RNA products were isolated, polyadenylated, and sequenced by direct RNA Oxford Nanopore sequencing (**Fig. 1B**). Direct RNA sequencing was used to reduce biases associated with classical short read RNA sequencing approaches including PCR amplification or the ability to detect short RNA species. The 3′ end of each transcript was mapped to the starting library to define the position of RNA polymerase II at the time of quenching (Fig. 1C). Variability in library preparation and transcript length were controlled for by adding an RNA spike-in consisting of five RNAs with random sequence compositions and lengths. The shortest unambiguously mapped RNA corresponds to 18 nucleotides (nt) (**Fig. S1 D, E** and **Methods**).

**Table 1:** Cryo-EM data collection, refinement, and validation statistics.

|  | Sidetracked RNA polymerase II elongation complex (EMD-XXXX) (PDB: XXXX) | Pretranslocated RNA polymerase II elongation complex (EMD-XXXX) (PDB: XXXX) |
| --- | --- | --- |
| **Data collection and processing** |  |  |
| Magnification | 105,000 | 105,000 |
| Voltage (kV) | 300 | 300 |
| Electron exposure (e-/Å<sup>2</sup>) | 53.8 | 53.8 |
| Defocus range (µm) | 0.25-2.25 | 0.25-2.25 |
| Pixel size (Å) | 0.822 | 0.822 |
| Symmetry imposed | C1 | C1 |
| Initial particle images (no.) | 2,546,600 | 2,546,600 |
| Final particle images (no.) | 36,583 | 23,701 |
| Map resolution (Å) | 2.9 | 2.8 |
| FSC threshold | 0.143 | 0.143 |
| Map resolution range (Å) | 2.9-10 | 2.8-10 |
| **Refinement** |  |  |
| Initial models used (PDB code) | 8A40 | 8A40 |
| Map sharpening B factor (Å<sup>2</sup>) | -20 | -20 |
| **Model composition** |  |  |
| Non-hydrogen atoms | 31939 | 31406 |
| Protein residues | 3792 | 3734 |
| Nucleotides | 80 | 76 |
| Ligands | 9 | 9 |
| **B factors (Å<sup>2</sup>)** |  |  |
| Protein | 158.93 | 160.82 |
| Nucleotide | 239.23 | 256.65 |
| Ligand | 237.54 | 246.57 |
| R.m.s. deviations |  |  |
| Bond lengths (Å) | 0.004 | 0.002 |
| Bond angles (°) | 0.537 | 0.501 |
| **Validation** |  |  |
| MolProbity score | 1.74 | 1.7 |
| Clashscore | 4.89 | 4.15 |
| Poor rotamers (%) | 1.11 | 1.27 |
| **Ramachandran plot** |  |  |
| Favored (%) | 92.9 | 93.55 |
| Allowed (%) | 7.1 | 6.45 |
| Disallowed (%) | 0 | 0 |

**Figure 1.**
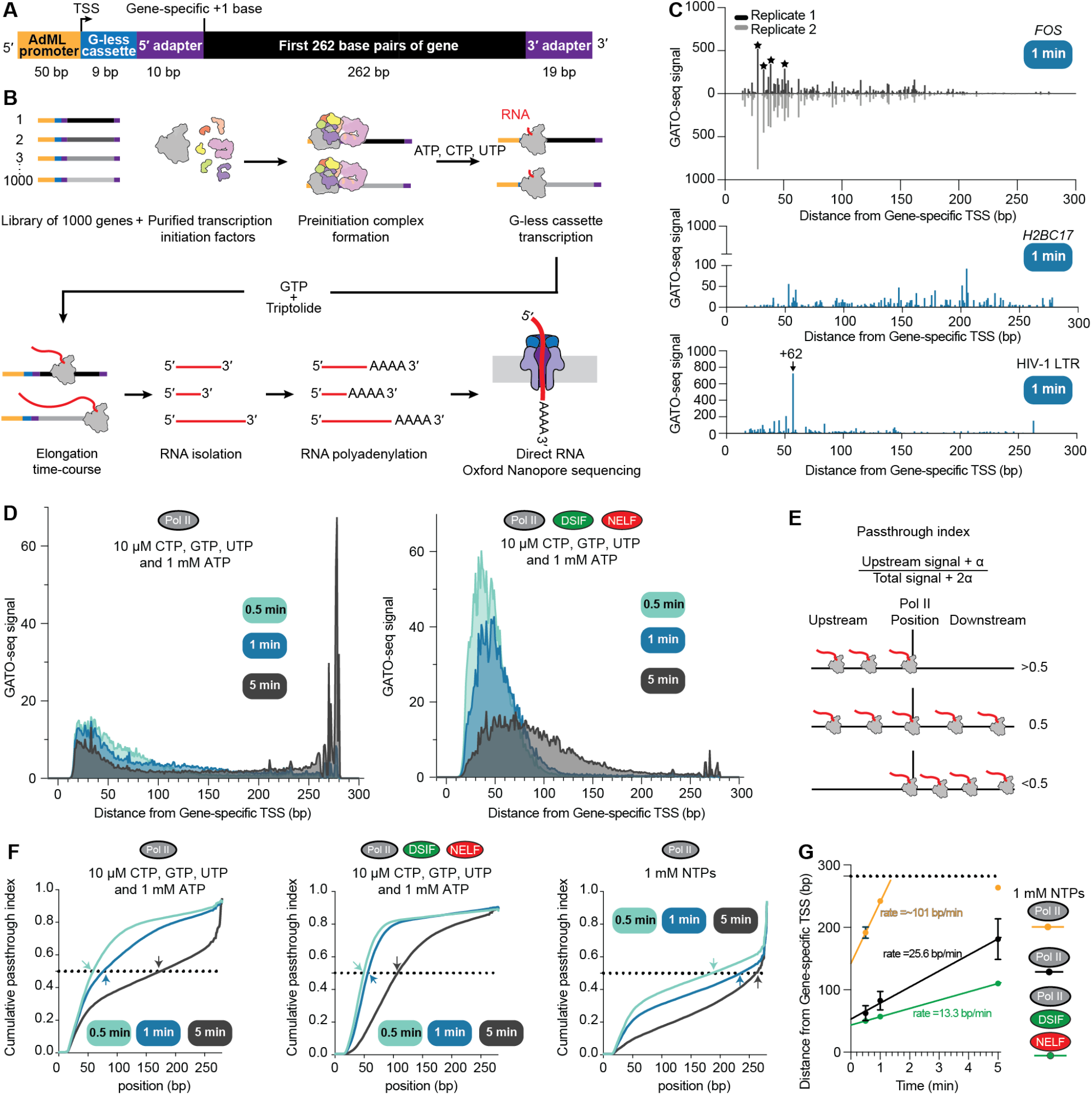
GATO-seq captures RNA polymerase II promoter proximal behavior. (**A**) Schematic of DNA template design employed in GATO-seq to study pausing. (**B**) Schematic of GATO-seq experimental design. (**C**) GATO-seq signal in gene-track representation for 3 genes. Top panel shows signal from two biological replicates, and bottom panels show the average signal from two biological replicates. (**D**) Metagene plots generated from the average signal across the sequence library at three different time points from two biological replicates employing 10 µM CTP, GTP, UTP and 1 mM ATP. Left, RNA polymerase II without additional factors and right with 3-fold molar excess of DSIF and NELF (**E**) Schematic of passthrough index estimation. (**F**) Passthrough index cumulative distributions at different time points from two biological replicates. Left, RNA polymerase II with 10 µM CTP, GTP, UTP and 1 mM ATP, center, with DSIF and NELF and right without additional factors and 1 mM NTPs. Arrows indicate cumulative distribution midpoint (y=0). RNA polymerase II: 60, 73 and 160 bp/min, DSIF-NELF: 48.5, 53.5 and 103.5 bp/min and 1 mM NTPs: 172, 228 and 256 bp/min) (**G**) Cumulative distribution midpoint plots plotted by time. Rates were estimated from a linear fit. Error bars represent the standard deviation from two biological replicates. Dotted line indicates the end of the template.

We first assessed whether GATO-seq could report on relative RNA polymerase II elongation velocity. Low NTP concentrations, (10 µM CTP, UTP, GTP and 1 mM ATP, required for TFIIH helicase and kinase activity), were employed to enhance the detection of paused RNA polymerase II complexes^54,55,58^. The normalized number of 3′ ends, termed GATO-seq signal intensity, defined the relative abundance of RNA polymerase II at each nucleotide position over time. Time course experiments performed in the absence of additional factors showed a steady progression of RNA polymerase II through the gene body where RNA 3′ ends were primarily observed within the first 100 bp after 30 s of elongation, extended beyond 100 bp and appeared at the end of the template after 1 minute (**Fig. 1D, Fig. S1F**). Fulllength transcripts were most abundant after 5 minutes (14.8 ± 1% of all 3′ ends) (**Fig. 1D**). When the time course was extended to 30 minutes, the amount of full-length transcript was not substantially different to that detected after 5 minutes (**Fig. S1G**). To estimate the relative velocity of RNA polymerase II, we calculated the ratio of upstream signal to total GATO-seq signal at each genomic position on a per gene basis (passthrough index) (**Fig. 1E**). The time dependent change in the cumulative distribution midpoint (y = 0.5) of the passthrough index was used to approximate the average elongation rate and corresponded to 25.6 ± 3.8 bp/min (**Fig. 1F, G**).

We next assessed whether the addition of pause associated protein complexes DSIF-NELF altered the relative elongation velocity of RNA polymerase II ^14–16,54,55,65^. Addition of DSIF-NELF decreased elongation rates by ∼2-fold (rate = 13.3 ± 0.43 bp/min) compared to the sample lacking DSIF-NELF (**Fig. 1D, F, G, Fig. S1H**). Finally, we tested if GATO-seq can distinguish differences in velocity produced by changes in NTP concentration. We specifically assessed whether 1 mM of NTPs, a concentration closer to physiological nucleotide levels, affected transcription elongation. Under these conditions, RNA polymerase II reached the end of most templates after 30 s. The cumulative density midpoint at 30 s and 1 minute were used to obtain a relative velocity of 101 ± 13 bp/min, which is likely an underestimate of the velocity (**Fig. 1F, G, Fig. S1I**).

### GATO-seq reports RNA polymerase II pausing behavior

We next designated positions in the library where RNA polymerase II shows pausing behavior. Pause sites are locations where RNA 3′ ends are significantly enriched compared to the rest of detected 3′ ends, and decay over time, indicating that RNA polymerase II has escaped the pause. In our data, 3′ peaks corresponding to previously established pause sites were observed, such as the HIV-1 Long Terminal Repeat +62 pause site and FOS ^54,59,66^ (**Fig. 1C**). To compare across time points, we first normalized the GATO-seq signal as fraction of total signal within each gene (**Fig. S2A**). Pause sites were then designated as positions where GATO-seq signal was ≥4 standard deviations above the gene signal average for at least two consecutive time points (**Fig. 2A, Fig. S2A**). As expected, more pause sites were observed at lower nucleotide concentrations than at higher concentrations (1702 versus 516 sites, respectively) (**Fig. 2B**). Similarly, addition of DSIF-NELF to low nucleotide containing samples resulted in the identification of 2718 pause sites (**Fig. 2B**). Subsets of pause sites overlapped across conditions. Specifically, 503 sites overlapped between low nucleotide samples lacking or con-taining DSIF-NELF. This overlap increased when the pause detection threshold was decreased from four to three or two standard deviations above the average (over-lap = 891 and 1253, respectively), suggesting that DSIF-NELF increase the pause propensity at otherwise weak pause sites (**Fig. 2C**). Between low and high nucleotide concentration samples, 128 positions overlapped (**Fig. 2C**).

**Figure 2.**
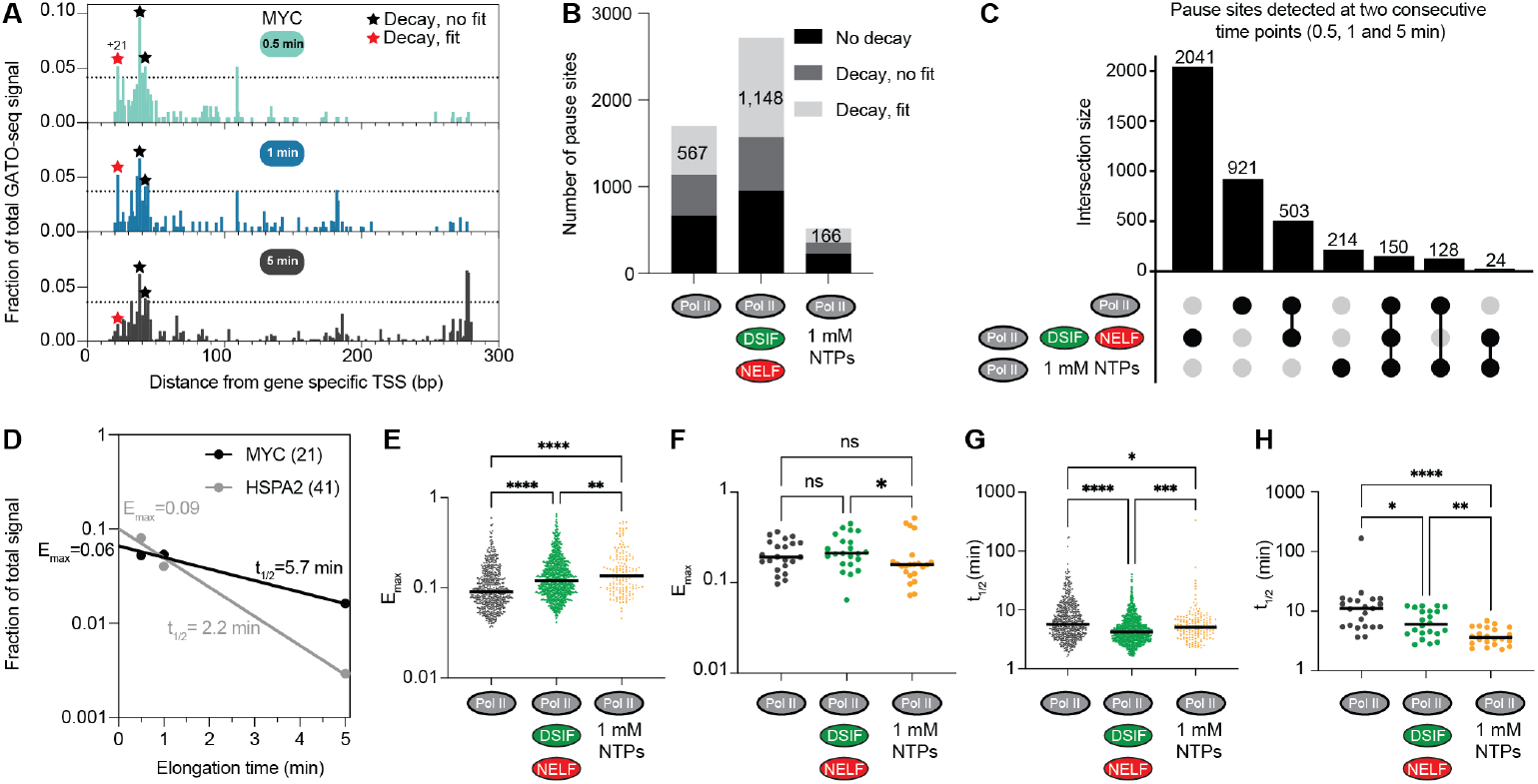
GATO-seq reports RNA polymerase II pausing behavior. (**A**) Representative normalized gene track for the MYC gene with RNA polymerase II and 10 µM CTP, GTP, UTP and 1 mM ATP. The RNA 3’end signal for each position along the gene is reported as its fraction of the total signal. Horizontal line indicates the pause detection threshold that is ≥4 standard deviations above the average signal. Stars indicate detected pause sites. The red star demarcates a pause site that displayed exponential decay shown in (D). Average of two biological replicates (**B**) Total number of pause sites detected in each experimental condition and categorized based on the decay behavior. (**C**) Intersection of pause site positions from three indicated experimental conditions displayed as an UpSet plot. (**D**) Fraction of GATO-seq signal versus time plotted for the +21 MYC and +41 HSPA2 pause sites. Both sites display exponential decay in GATO-seq signal over time. Points are average values from two biological replicates. E_max_ and t_1/2_ values are indicated. (**E**) Distribution of E_max_ values from pause sites displaying exponential decay in (B) across the three indicated experimental conditions. Statistical significance was assessed using a nonparametric Kruskal-Wallis test (p-values *=0.032 **=0.0021, ***<0.0002 and ****<0.0001). Subset of E_max_ values shared across the three experimental conditions, displayed as in (E). (**G**) Distribution of t_1/2_ from indicated experimental conditions. Statistical comparison as in (E). (**H**) Subset of t_1/2_ values shared across the three experimental conditions, displayed as in (G).

We next quantified pausing behavior using two previously defined empirical values^60,61^. These values are calculated by plotting the fraction of total signal for each pause position versus time. The maximal pause efficiency (E_max_) is proportional to the maximal fraction of RNA polymerase II that pauses at a particular pause position and is defined by the y-intercept^60,61^ (**Fig. 2D**). The pause half-life (t_1/2_) is calculated as the decay of signal over time. These plots were only analyzed for pause sites where both the GATO-seq signal decayed over time (*i*.*e*., the lowest signal was observed at the 5-min timepoint) (**Fig. 2B**) and robustly fit to an exponential decay model (**Fig. 2B, D**). These plots revealed a variety of behaviors at each pause site across conditions. We note that fitting only 3 time points can lead to over and underestimation of t_1/2_. We thus evaluated how using 3 or 5 time points from a 30-minute time course experiment affected our fit of t_1/2_. Under these conditions, t_1/2_ at 69 % of pause sites was not substantially different (± 25% value) between 3 or 5 time point analyses. Conversely, 16 % of pause sites showed underes-timation and 15% of pause sites showed overestimation of t_1/2_ when only using 3 time points for fitting (**Fig. S2B**).

The E_max_ corresponds to how readily RNA polymerase II enters a paused state at a particular location. As expected, samples containing DSIF-NELF had higher median E_max_ values than matched samples lacking DSIF-NELF (median 0.119 and 0.089, respectively) (**Fig. 2E**). Higher NTP concentrations also resulted in an increase of the median E_max_ value (0.135) (**Fig. 2E**). This is likely because fewer pause sites are called in the higher nucleotide condition, and the detected pauses are strong pause sites. Indeed, lowering the pause detection threshold to include pauses that were only three or two standard deviations above the average, resulted in lower median E_max_ (0.100 and 0.072, respectively) (**Fig. S2C**). Consistently, the opposite effect was ob-served for DSIF-NELF condition, where the median E_max_ values increased to 0.143 and 0.165 as the pause detection threshold increased to five and six standard deviations, respectively (**Fig. S2C**). We next examined the median E_max_ values for sites that were shared among all three conditions (22 sites). At these sites, the E_max_ median is elevated for Pol II-DSIF-NELF versus Pol II samples (0.211 versus 0.192, respectively) whereas a reduction in the median E_max_ is observed for the 1 mM NTP containing sample (0.158) (**Fig. 2F**).

The t_1/2_ measures the half-life of RNA polymerase II at a pause site. Unexpectedly, addition of DSIF-NELF decreased the median t_1/2_ compared to a matched sample lacking DSIF-NELF (t_1/2_ = 4.24 versus 5.71 minutes, respectively) (**Fig. 2G**). The 1 mM NTP concentration sample had a median t_1/2_ of 5.04 minutes (**Fig. 2G**). Unlike the median E_max_, median t_1/2_ values were largely independent of the pause detection threshold used (**Fig. S2D**). We next compared the median t_1/2_ for pause sites that were preserved across conditions. We observed an overall reduction in median t_1/2_ when DSIF-NELF were present or when physiological nucleotide concentrations were employed (11.04, 6.02 and 3.62 min, respectively) (**Fig. 2H**).

In summary, DSIF-NELF reduce both RNA polymerase II velocity and the dwell time of RNA polymerase II at pause sites, whereas physiological nucleotide concentration drastically reduces the number of pause sites detected.

### GATO-seq detects TFIIS specific effects on pause site selection

RNA polymerase pausing is often associated with “offline” active site states that are not competent for nucleotide incorporation. These include RNA-DNA hybrid translocation intermediates known as half-translocated or tilted states and reverse translocation events termed backtracking^47,54,67,68^. TFIIS is a factor that can overcome these offline states by realigning the RNA-DNA hybrid and stimulating endonucleolytic cleavage of backtracked RNA by RNA polymerase II^47,49,67,69–73^. We thus assessed whether pause positions changed upon the addition of TFIIS. TFIIS was added to reactions with GTP and triptolide and reactions were stopped after 1 and 5 minutes (**Fig. 3A, B, C**).

**Figure 3.**
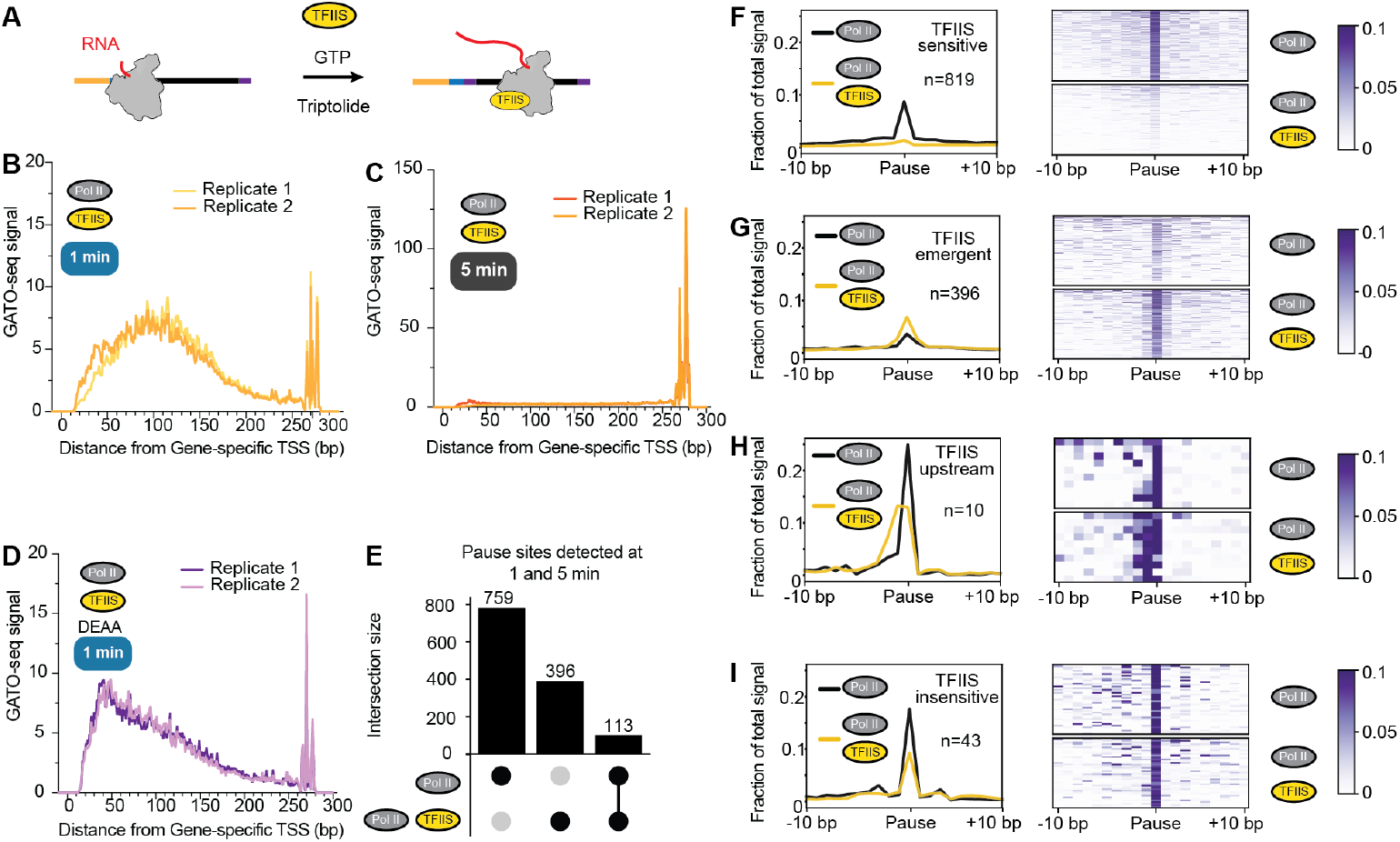
TFIIS specific effects on pause site selection. (**A**)Schematic of TFIIS addition to GATO-seq experiments (**B-D**) Metagene plots of the average GATO-seq signal across the sequence library. GATO-seq signal in the presence of TFIIS for (**B**) 1 minute and (**C**) 5 minutes, or (**D**) TFIIS (DEAA) for 1 minute. RNA polymerase II:TFIIS molar ratio is 1:0.2. Experiments were performed with 10 µM CTP, GTP, UTP and 1 mM ATP. Signal from two biological replicates. (**E**) UpSet plot showing the intersection of pause sites found with and without TFIIS in both 1-and 5-minute time points. (**F-I**) Profile (left) and heatmap (right) of pause sites pileups from 20 bp windows with the pause site centered at the 10_th_ position. Average fraction of signal from two biological replicates. Sites are categorized based on the effect of TFIIS. (**F**) TFIIS sensitive (**G**) TFIIS emergent (**H**)TFIIS upstream shift (**I**)TFIIS resistant.

TFIIS addition resulted in a cumulative density midpoint of 120 ± 2.8 bp after 1 minute of transcription, which is ∼38 bp further downstream than RNA polymerase II without TFIIS (**Fig. 3B, Fig. S2E**) and led to a nearly 2-fold increase of full-length transcripts after 5 minutes of transcription compared to samples lacking TFIIS (29 ± 6% versus 14.8 ± 1%, respectively, of all detected transcripts are full-length) (**Fig. 3C**). TFIIS DE282-283AA is a mutant that cannot stabilize second-metal binding in the RNA polymerase II active site and therefore does not support RNA cleavage by RNA polymerase II^49,70,72,74,75^. Unexpectedly, inclusion of TFIIS DE282-283AA resulted in a cumulative density midpoint of 97 ± 1.4 bp after 1 minute of transcription, which is ∼15 bp downstream of RNA polymerase II without TFIIS (**Fig. 3D, Fig. S2E**), suggesting that it may positively impact transcription elongation in a mechanism that does not involve second metal binding. To define the effect of TFIIS at specific pause sites, we called pause sites at locations where the fraction of total GATO-seq signal per gene was ≥4 standard deviations above the average and present at both 1 and 5 minutes. Pause positions between samples lacking or containing TFIIS were then compared. A total of 873 and 510 unique pause sites were detected in the absence and presence of TFIIS, respectively (**Fig. 3E**). We then compared pause positions within ±3 nt between samples lacking TFIIS to those containing TFIIS. Most pause sites were only detected in the absence of TFIIS (759 sites, 59.8 % of all pause sites) indicating that these pause sites are sensitive to TFIIS (TFIIS sensitive) (**Fig. 3E, F**). Some pause sites were only detected when TFIIS was present (396 sites, 31.2 % of total) (**Fig. 3E, G**). Notably, a subset of these sites (55 sites) showed elevated GATO-seq signal in the absence of TFIIS compared to the surrounding sites but did not meet the criteria to be called. These additional pause sites may be a consequence of RNA polymerase II being more efficiently released from upstream pause sites. Supporting this possibility, the median passthrough index at these pause sites after 1 minute of transcription was 0.5 in the absence of TFIIS, and 0.29 with TFIIS (**Fig. S2F**), indicating higher RNA polymerase II upstream occupancy in the absence of TFIIS. Finally, 113 (8.9 %) sites were shared between the two conditions (**Fig. 3E, H, I**). Shared pause sites display three different behaviors. Cluster 1 pauses displayed an average ∼50% decrease in signal upon TFIIS treatment, indicating sensitivity to TFIIS and therefore, we classify them as such (**Fig. 3F**). Cluster 2 showed a 1-2 bp upstream shift upon TFIIS addition (TFIIS upstream sites) (**Fig. 3H**), and cluster 3 showed minimal changes in GATO-seq signal and position upon TFIIS addition (TFIIS insensitive) (**Fig. 3I**). Of these 43 sites, 19 were also found in the 1 mM NTP condition, whereas 29 were found in reactions with DSIF-NELF, further indicating they are strong pause positions. These results show that most pauses were overcome by TFIIS addition; however, a subset of pauses were insensitive to TFIIS.

### GATO-seq uncovers consensus “super pause” sequence

We next defined sequence features of the pause sites that were insensitive to TFIIS addition. These sequences were aligned relative to the pause site, and a sequence logo was generated (**Fig. 4A**). TFIIS insensitive pauses exhibited a conserved upstream sequence composition, including a purine (R)-rich stretch from positions -13 to -7, harboring an enriched guanosine at position -8, a cytosine or adenosine (M) at position -10, a cytosine at position -2, a pyrimidine (Y) at the 3′ end (-1), an incoming guanosine (+1) and +2, +3 cytosines (consensus R_-13_R_-12_R_-11_M_-10_R_-9_G_-8_A_-7_C_-2_Y_-1_G_+1_C_+2_ C_+3_) (**Fig. 4A**). This motif shows similarities and differences to previously reported pause motifs (**Fig. S1A**).

**Figure 4.**
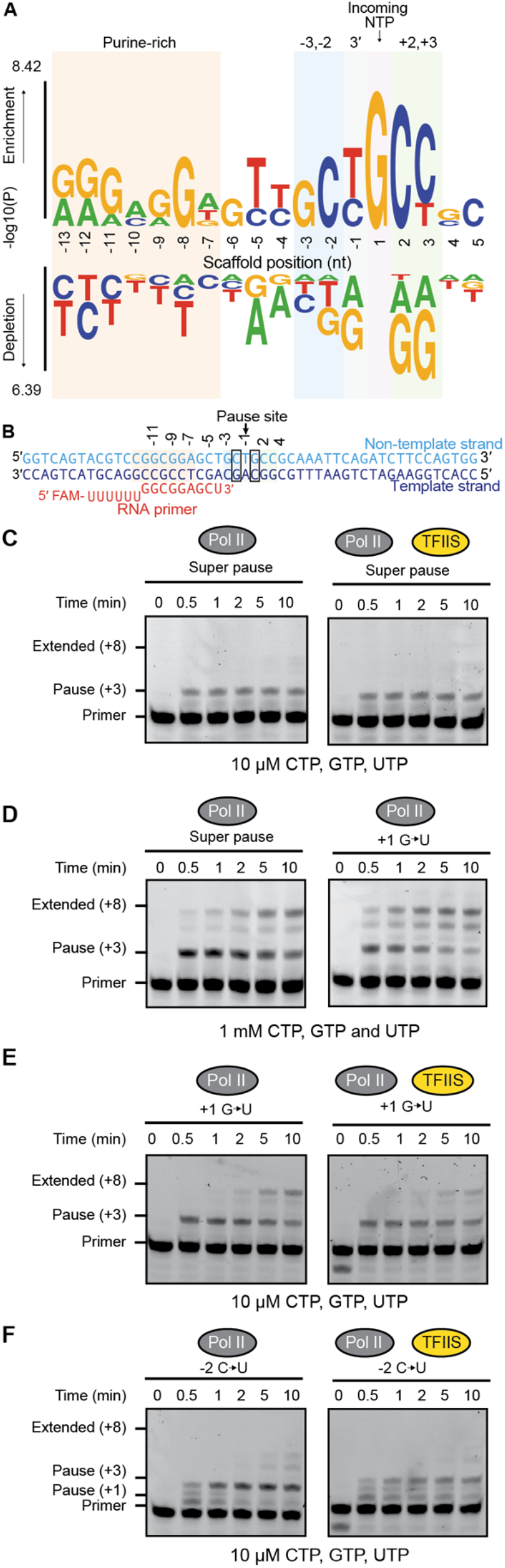
GATO-seq uncovers “super pause” consensus sequence. (**A**) Sequence logo derived from TFIIS insensitive sites (n=43 sequences). (**B**) Super pause consensus sequence scaffold used for RNA extension assays (**C**) RNA primer extension time course experiments in the absence or presence of TFIIS and 10 µM CTP, GTP and UTP. RNA polymerase II:TFIIS used at a molar ratio of 1:0.2. All experiments were performed at least three times on super pause consensus scaffold (**D**) RNA extension assay with 1 mM CTP, GTP and UTP on the super pause scaffold and the super pause scaffold with a +1G to U substitution. (**E**) RNA extension assay as in (C) with super pause scaffold caring a +1G to U substitution (**F**) RNA extension assay as in (C) caring a -2C to U substitution.

We designed a consensus pause sequence using the features detected at TFIIS insensitive pause sites (**Fig. 4B**). RNA extension assays were first executed on this consensus sequence with 10 µM CTP, UTP, and GTP thereby permitting extension up to +8. The pause site was placed at position +3. RNA extension reactions recapitulated the observations of GATO-seq (**Fig. 4C**), showing robust RNA polymerase II pausing with minimal extension from the pause site even after ten minutes, and no pronounced change in pause position upon TFIIS inclusion (**Fig. 4C**). Similarly, RNA extension assays conducted with 1 mM of each NTP (minus ATP) showed the pause band remained the most prominent feature throughout a 10-minute time course (**Fig. 4D, Fig. S3A**). In contrast, a sequence lacking the enriched consensus pause elements referred to as the anti-pause sequence showed no pausing behavior at either nucleotide concentration (**Fig. S3B**). We next tested the role of invariant sequence features in the consensus motif on RNA polymerase II pausing behavior. An incoming guanosine is strongly associated with TFIIS insensitive pausing behavior. Substitution of this guanosine to uracil resulted in full-length product detection two minutes after nucleotide addition and more efficient pause release with 1 mM NTPs (**Fig. 4D, E, S3C**). Conversely, substitution of the -2 cytidine with uracil resulted in sensitivity to TFIIS and the appearance of a 2-nt shorter product (**Fig. 4F, S3D**). The -2 cytidine is not enriched in the TFIIS sensitive consensus sequence (**Fig. S3E**). This indicates that specific features of the consensus sequence confer different functions on RNA polymerase II pausing behavior. Together, these experiments demonstrate that the TFIIS insensitive consensus pause sequence induces a very stable and strong pause that is unaffected by TFIIS, lasts for an extended period (>10 minutes) and is minimally affected by higher nucleotide concentrations. We thus refer to this consensus sequence as a “super pause” sequence.

### RNA polymerase II does not prematurely terminate at the super pause

RNA polymerase II can terminate or arrest at strong pause sites^76–78^. To investigate whether RNA polymerase II is engaged with DNA on the super pause consensus sequence, we first conducted pyrophosphorolysis experi-ments^79,80^. High concentrations of pyrophosphate induce RNA polymerase II to reverse bond formation at the terminal nucleotide. This can be detected as a shortening of the 3′ RNA end that is only possible when RNA polymerase II is engaged with the DNA (**Fig. 5A**). Addition of pyrophosphate to samples containing RNA polymerase II at the consensus super pause site induced strong pyrophosphorolysis indicating no premature termination (**Fig. 5A**).

**Figure 5.**
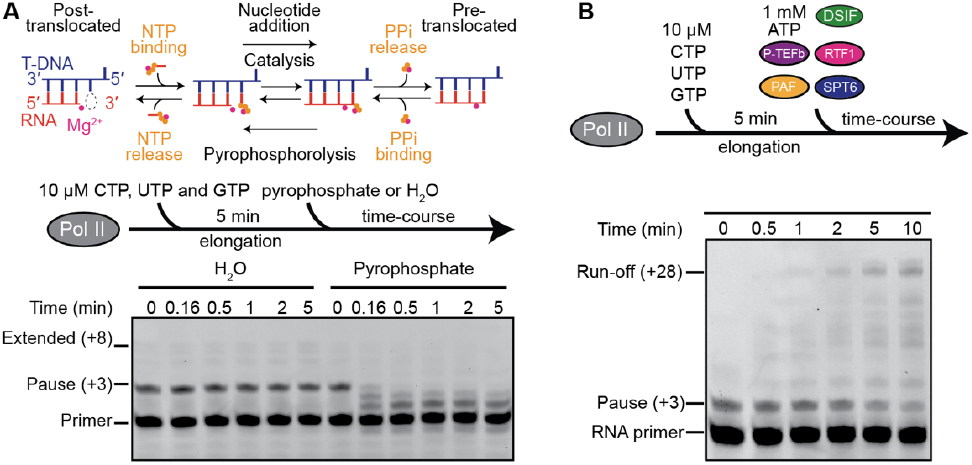
RNA polymerase II does not prematurely terminate at the super pause. (**A**) RNA pyrophosphorolysis time course experiments. Top panel: schematic of reversible catalysis of nucleotide addition and pyrophosphorolysis carried by RNA polymerase II. Bottom panel: experimental schematic and representative gel of an experiment performed after 5 minutes of elongation on the super pause consensus scaffold using 10 µM NTPs, followed by the addition of H_2_O or 1 mM pyrophosphate. All experiments were performed at least three times. (**B**) RNA primer extension time course on the super pause consensus scaffold after 5 minutes of elongation using 10 µM NTPs followed by the formation of EC* (1 mM ATP, and 1:3 molar ratio of RNA polymerase II: DSIF, P-TEFb, PAF1c (with RTF1) and SPT6). Addition of ATP in this experiment leads to complete primer extension (Run-off +28 band). All experiments were performed at least three times.

Next, we investigated whether RNA polymerase II is irreversibly arrested at the super pause site. Specifically, the activated RNA polymerase II complex (EC*, RNA polymerase II with elongation factors DSIF, PAF1c, RTF1, and SPT6, facilitated by the kinase P-TEFb) can release reversibly paused RNA polymerase II from pause sites^22,23,28^. EC* was formed before or after RNA polymerase II reached the super pause site (**Fig. 5B, Fig. S4A, B**). Fulllength products were clearly detectable at 30 seconds and 1 minute respectively, using 10 µM CTP, GTP, UTP and 1 mM ATP (required for P-TEFb kinase activity), indicating that RNA polymerase II at the super pause is not arrested and can be converted into a processive enzyme. Importantly, EC* activation could also be triggered after five minutes when the RNA polymerase II–DSIF–NELF complex was paused at the super pause site (**Fig. S4C**). Together, these findings indicate that RNA polymerase II paused at the super pause site is in a stable but largely reversible offline state.

### Cryo-EM structure of RNA polymerase II at the super pause

Prior structures of paused RNA polymerase complexes have revealed backtracked and tilted RNA-DNA hybrid active site conformations^39,54,55,69,70,81^. These offline conformations can be alleviated by TFIIS addition. To understand the molecular basis for RNA polymerase II pausing at the super pause consensus sequence and its insensitivity to TFIIS addition, we visualized RNA polymerase II on the super pause consensus sequence using single-particle cryo-EM. Complexes for cryo-EM were prepared on the super pause consensus sequence and RNA polymerase II was permitted to transcribe for 5 minutes. The actively transcribing complexes were then purified and mildly crosslinked before preparing cryo-EM grids (**Fig. S5A-C**). We collected 2,546,600 particles of which 862,521 showed high resolution features for RNA polymerase II (**Fig. S5D, E**). Peripheral resolutions ranged from ∼6-10 Å, whereas the resolution within the active site was ∼2.9 Å, and allowed for unambiguous assignment of the corresponding bases in the DNA-RNA hybrid (**Fig. S6 and S7 A-F**).

Classification yielded three active site conformations, including pretranslocated (23,701 particles), posttranslocated (24,149 particles), and a previously unobserved backtracked state (36,583 particles) hereafter referred to as “sidetracked” (**Fig. 6A, B, S7A-F, Fig. S8**). Initial models were built for the pretranslocated and side-tracked states using published RNA polymerase II elongation complexes structures^81^, and the nucleic acid sequence and stereochemistry were manually adjusted and refined (**Table 1**). The posttranslocated state showed heterogenous sequence composition in the RNA polymerase II active site suggesting a mixture of states and was thus not modelled (**Fig. S7F**). The pretranslocated and sidetracked states have transcribed three nucleotides (**Fig. S7G**). All classes exhibit a mobile trigger loop conformation. The sidetracked state is incompatible with trigger loop folding, an event that is required for nucleotide addition and is generally permitted in singly backtracked states^82–86^ (**Fig. 6C**). Finally, structures of mammalian RNA polymerase II-TFIIS complexes show high flexibility for TFIIS domain III, the domain that is responsible for TFIIS function, preventing appropriate modelling of the domain^39,55,81,87^. To overcome this limitation, we overlaid our structures with the structure of TFIIS from a yeast RNA polymerase II-TFIIS rescue intermediate where domain III is well resolved^69^. This shows that the sidetracked base would clash with the positioning of TFIIS domain III (**Fig. 6D**). In contrast, TFIIS domain III is readily accommodated in the pretranslocated state, a state where TFIIS is generally not thought to act^88^ (**Fig. S9A**). This provides a structural basis for the lack of evident TFIIS action on the super pause sequence (**Fig. 4C**).

**Figure 6.**
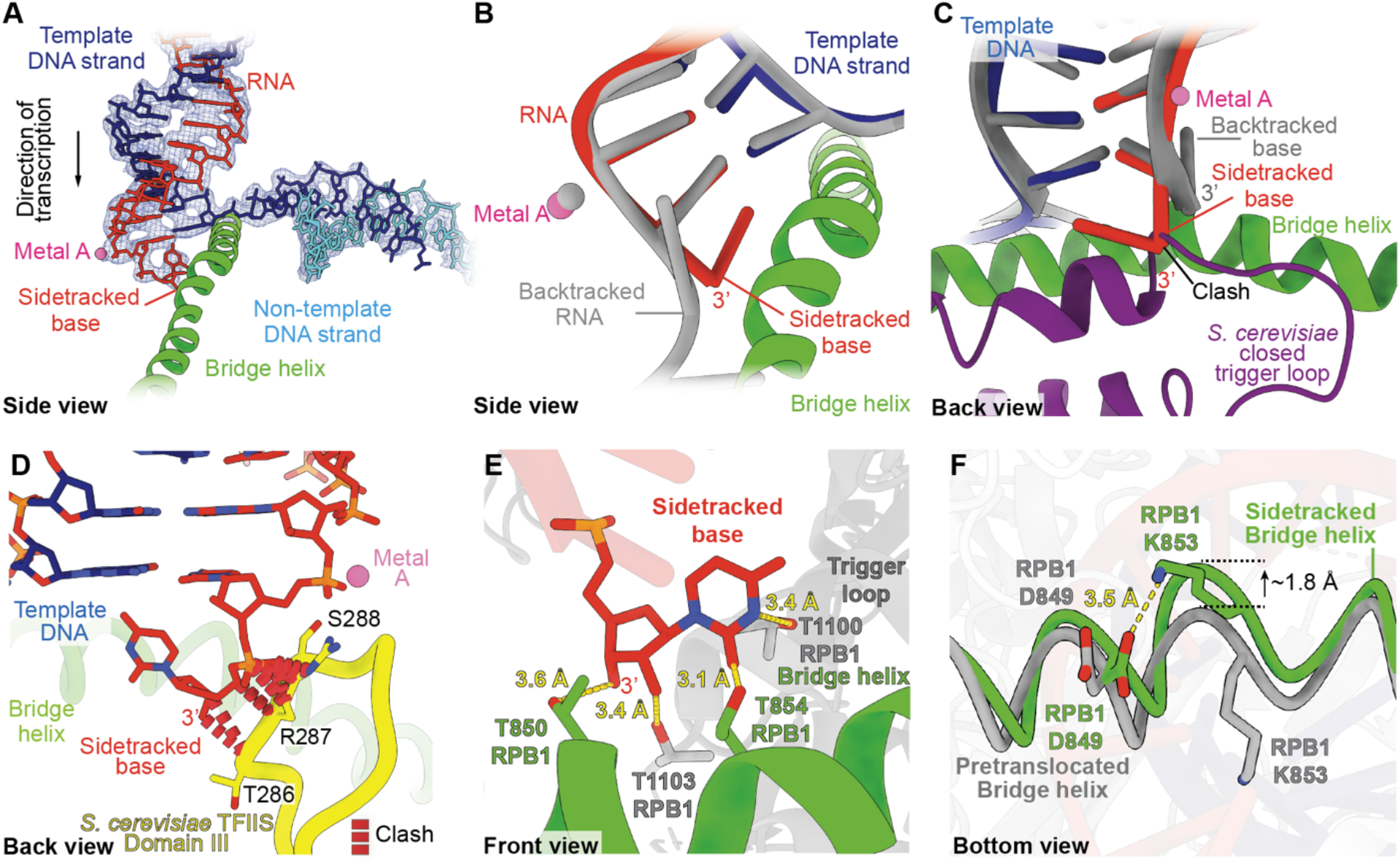
Cryo-EM structure of RNA polymerase II on the super pause consensus sequence. (**A**) Cryo-EM density (blue mesh) of template DNA (blue)-RNA (red) hybrid in the sidetracked state. Bridge helix is shown in green and metal A in magenta. (**B**) Comparison of sidetracked register (RNA in red, template-DNA strand in blue) to a 6 nt backtracked RNA shown in gray (PDB 8A3Y). Bridge helix from sidetracked structure shown in green (**C**) Back view, overlay of sidetracked base (red), 1-nt S. cerevisiae backtracked RNA-DNA hybrid (gray) (PDB 3GTG) and S. cerevisiae closed trigger loop (purple) (PDB 2E2H). Steric clash between closed trigger loop and sidetracked base is indicated with a black arrow. Bridge helix from sidetracked structure is shown in green and metal A in magenta (**D**) Overlay of sidetracked RNA (red) and S. cerevisiae TFIIS Domain III (PDB 3PO3) shown in yellow. Red dashed lines indicate a steric clash between the sidetracked base and TFIIS residues shown as sticks. Bridge helix shown in green and metal A in magenta (**E**) Front view, RNA polymerase II threonine pocket. Threonine side chains and sidetracked RNA nucleotide shown as sticks. (**F**) Comparison of RNA polymerase II bridge helix in the sidetracked (green) and pretranslocated (gray) states on the super pause consensus sequence. Bridge helix residues RPB1 D849 and R853 shown as sticks.

The most prominent feature of the sidetracked state is the position of the RNA 3′ end. Previously determined structures of backtracked RNA polymerases show an RNA end that is bent by ∼120º relative to A-form RNA^69,70^. In the sidetracked state, the RNA was projected in the opposite orientation and was bent by ∼50° (**Fig. 6B, Fig. S9B, C**)^39,68–70,81^. This positioning would only allow for the back-tracking of a single base, whereas canonical backtracking can involve the extrusion of multiple RNA bases through the pore and funnel region (**Fig. 6B, Fig. S9D-H**). Additionally, the penultimate RNA base in the sidetracked conformation formed a canonical base pair with the DNA template strand, and the hybrid remained planar as observed in the pretranslocated state (**Fig. S9J, K**).

The sidetracked base forms several key interactions with RNA polymerase II. Specifically, RPB1 bridge-helix residues T850, T854, and trigger loop N-terminal helix residues T1100 and T1103 form a “threonine pocket” to coordinate the ribose and pyrimidine rings (**Fig. 6E, S9L**). The threonine pocket appears to be conserved among multi-subunit bacterial and eukaryotic RNA polymerases (**Fig. S9M**). RPB1 N493 contacted the α-phosphate of the terminal RNA base. These interactions positioned the pyrimidine ring >13 Å from the RPB2 gating Y724 and RPB2 fork residue E516, which typically stabilize the first canonically backtracked RNA base (**Fig. S9B**)^70^. The threonine pocket appears to more readily accommodate pyrimidines over purines, and purine bases, specifically guanosine, may clash with the pocket (**Fig. S9N**). This structural observation is consistent with the depletion of purines in the super pause consensus sequence at the 3’ end (**Fig. 4A**). The formation of the thre-onine pocket requires bending of the bridge helix by ∼2 Å relative to the conformation adopted in the pretranslocated state (**Fig. 6F**). This bending was stabilized by a salt bridge formed between RPB1 D849 and K853, as has been observed in conventional backtracking structures of RNA polymerase II and bacterial RNA polymerase (**Fig. S9F-H**)^68,69,73^. Molecular dynamics simulations have suggested that RPB1 bridge helix residue T854 (yeast T831) acts as a sensor for appropriate base pairing between the last incorporated RNA base and the DNA base at the site of nucleotide incorporation^89^. This sensor activity may allow for bridge helix bending when inappropriate base pairing is detected to allow for fraying and extrusion of the misincorporated RNA base out of the pore region (**Fig. S9I**). In addition to this role, our structure showed that T854 appears to directly support the sidetracked state.

Our structural data obtained from actively transcribing RNA polymerase II complexes show that RNA polymerase II is primarily in the sidetracked or pretranslocated state. This is notable for two reasons. The first is that this supports the idea that sidetracking is a reversible offline state where RNA polymerase II can cycle between the online pretranslocated state and the offline sidetracked state. Second, previous work with bacterial RNA polymerase has shown that specific sequences at the site of nucleotide addition and in the incoming base favor pretranslocated states. The super pause sequence contains a terminal uracil and incoming guanosine. A terminal uracil is the base most likely to favor the pretranslocated state and an incoming guanosine can impede forward translocation by bacterial RNA polymerase^53,79,90^. In summary, our structural work shows that the super pause consensus sequence results in sidetracking, a novel form of backtracking that likely favors pyrimidines, and provides a molecular basis for understanding why the super pause is incompatible with TFIIS domain III engagement.

### RNA polymerase II occupancy is enriched at super pause sites in cells

Finally, we assessed how frequently RNA polymerase II accumulates at super pause sequences in human protein coding genes. We initially scanned the first 300 base pairs of 10,000 expressed protein-coding genes in HEK293T cells for the 43 experimentally derived super pause sequences. This analysis identified 4,455 hypothetical super pause sites distributed across 2,227 genes (**Fig. 7A**) (**Supplemental Table 1**). We generated background sequences with matched dinucleotide composition to the super pause sequences and found 1,819 sites (**Fig. 7A**). The distribution of hypothetical super pause sequences relative to the TSS, shows a mild enrichment within the first 50 bp of gene bodies compared to background sequences (**Fig. S10A**). We used these loci to compare the signal of RNA polymerase II occupancy on genes with hypothetical super pause sites to background sites using published chromatin immunoprecipitation coupled to exonuclease treatment (ChIP-exo) and PRO-seq data^91,92^ (**Fig. 7B, C**). Datasets collected from HEK293T cells where anti-RPB1 ChIP-exo was performed showed a more focused signal neighboring the TSS and lower signal further downstream into the gene body for hypothetical super pause sites, compared to the background sites (**Fig. 7B**). This behavior is consistent with prior reports indicating that promoter-proximal pausing hampers new initiation events, which may be the case in genes harboring hypothetical super pause sequences, leading to higher steady-state RNA polymerase II occupancy on genes without strong pausing^48,93^. Consistently, PRO-seq promoter-proximal signal at genes harboring hypothetical super pause sites was ∼30% higher than that of background sequences (**Fig. 7C**), indicating an elevated amount of engaged RNA polymerase II at promoter proximal regions with hypothetical super pauses. PRO-seq relies on the incorporation of a biotinylated NTP. Given the strength of the super pause sequence detected in our GATO-seq and RNA extension assays, it is likely that PRO-seq may not accurately inform on RNA polymerase II occupancy at these sites. Alternatively, NET-seq does not rely on modified nucleotide incorporation and could potentially capture sidetracked complexes. We sought to compare previously reported high-sensitive NET-seq signal at the hypothetical super pause to background sequences; however, the coverage was lower than PRO-seq and it was thus not possible to detect signal differences^94^ (**Fig. S10B**). GO analysis of hypothetical super pauses revealed enrichment of processes including response to stimulus and development (**Fig. S10C**). Background sequences did not show a significant enrichment of any functional class. In summary, the presence of super pause sites in cells could lead to enhanced accumulation or pausing of RNA polymerase II in promoter proximal gene regions.

**Figure 7.**
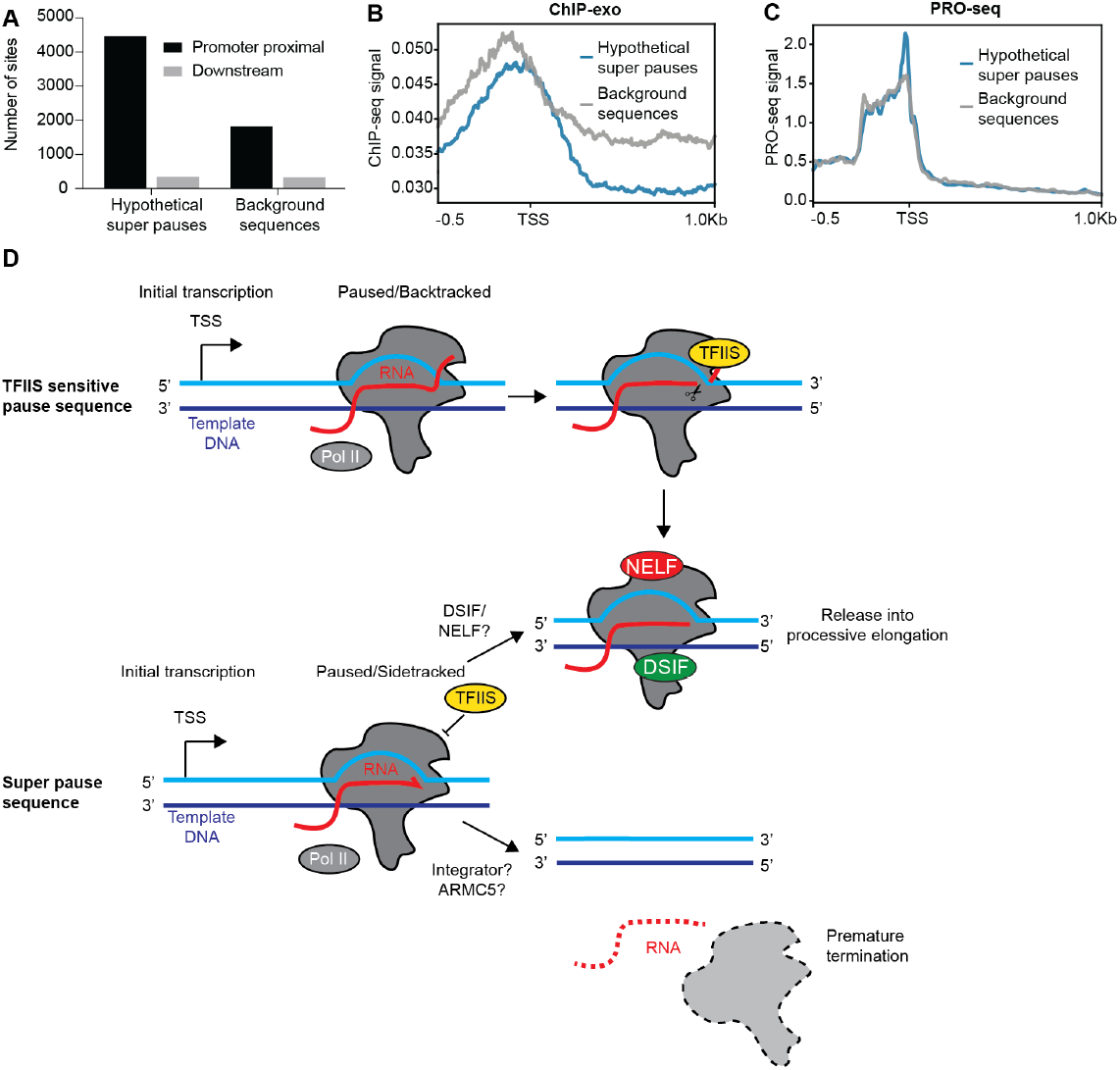
RNA polymerase II occupancy is enriched at super pause sites in cells. (**A**) Number of hypothetical super pause sequences and background sequences found within +1 to +300 (promoter proximal) and within +1000 and +1300 (gene body) windows on 10,000 protein coding genes. (**B and C**) Profile of genes with hypothetical super pause sites and background sites from (**B**) ChIP-exo and (**C**) PRO-seq. (**D**) Model of the role of RNA polymerase II sidetracking at super pause sites. During early elongation, RNA polymerase II can backtrack at TFIIS sensitive sequences, where TFIIS can help overcome non-productive states and RNA polymerase II can resume transcription. At super pause sequences, RNA polymerase II sidetracks thereby becoming resistant to TFIIS. Sidetracked complexes may associate with DSIF and NELF and/or cleared by termination factors, preventing partially assembled complexes to proceed into the gene body.

## Discussion

Here we have developed a reconstituted transcription platform, GATO-seq, to investigate RNA polymerase activity at scale. This assay in principle can be used on any DNA sequence of interest, monitor how transcription initiation or elongation behavior is altered by sequence, nucleotide concentration, or factor addition, and incorporate factors that compact and organize DNA. GATO-seq uses direct RNA sequencing and thus can report on single transcripts and is not biased by PCR amplification. As a proof of principle, we used GATO-seq to study mammalian RNA polymerase II elongation on a library composed of human promoter-proximal sequences.

### Sequence encoded RNA polymerase II offline states

RNA polymerases across all domains of life are subject to pausing. The active site of multi-subunit RNA polymerases is highly conserved suggesting that the fundamental mechanism of transcriptional pausing shares a common mechanistic basis^95^. DNA sequence has long been recognized as a determinant of RNA polymerase pausing, influencing both the position and dwell time of pausing. Several sequence features associated with pausing are conserved from bacteria to humans, including a strong DNA-RNA hybrid and a G-rich upstream fork junction where the template and non-template DNA strands reanneal (approximately -10 to -12 relative to the RNA 3′ end)^45,52,53,96^. In addition, the sequence surrounding the RNA 3′ end itself has been linked to prolonged pausing^52,59^. The super pause consensus sequence shares these sequence features. The super pause consensus is distinguished by strong conservation of two cytidines at positions +2 and +3, which are poorly enriched in the bacterial elemental pause consensus sequence. This feature is particularly notable because the super pause consensus closely resembles the HIV-1 LTR +62 pause sequence, including U_-1_G_+1_C_+2_. Mutations of the HIV-1 LTR +62 pause sequence that introduce C_+3_ increase pause duration^59,60^.

Cell-based approaches have defined consensus pause sequences which share some features with the super pause consensus sequence. Some studies have reported a preference for a terminal guanosine and an incoming pyrimidine^45,46^, whereas others have suggested a terminal cytidine and a weak preference for an incoming guanosine, similar to the super pause consensus sequence. G-rich fork junction features, however, are poorly enriched in all of these consensus sequences^48,51,97^ (**Fig. S1A**). Using GATO-seq, we were able to classify super pause sequences from a broader set of sequences that induce weaker pausing. Such sequence-level resolution is difficult to achieve in cell-based assays, where numerous additional factors, beyond TFIIS, DSIF, and NELF, contribute to pause-site selection and stability. Additionally, analysis of gene-specific transcription regulation in cell-based experiments is restricted to the genes that are expressed in a particular cell-line, and the depth of 3’end sequencing can be limiting. These complexities likely underlie the discrepancies between the super pause consensus identified here and pause consensus sequences inferred from cell-based approaches.

Our structural analysis of actively transcribing RNA polymerase II on the consensus super pause sequence revealed an offline active state conformation, termed side-tracked, that to our knowledge has not been previously reported. The sidetracked state is supported by a highly conserved threonine pocket formed by the bridge helix and trigger loop and appears to be selective for RNAs containing terminal pyrimidines. These results suggest that RNA polymerase II can adopt a sequence-dependent offline active site state that is resistant to pause rescuing factor TFIIS. Finally, the residues forming the threonine pocket are conserved across all domains of life and across human RNA Polymer-ases I, II, and III suggesting that sidetracking could be supported in other multisubunit RNA polymerases.

We speculate that sidetracked RNA polymerase II may be selectively removed from gene bodies by termination/attenuation to prevent aberrant transcription. This could explain why super pause complexes are poorly captured by cell-based assays. PRO-seq may be particularly insensitive to these states, as it requires additional nucleotide incorporation during the run-on step, which may be extremely slow or incompatible for sidetracked (and backtracked) complexes. Consistent with this idea, super pause-containing genes show only a modest promoter-proximal enrichment in PRO-seq datasets.

### The role of elongation factors at pause sites

In cells, DSIF-NELF associate with RNA polymerase II shortly after promoter escape. Although DSIF-NELF have been associated with increased pause duration, our results show that they slow down the overall rate of RNA polymerase II translocation and surprisingly decrease pause duration^58^. We speculate that DSIF-NELF may disfavor RNA polymerase II from entering some offline states, including backtracking, allowing for more rapid conversion into activated transcription elongation complexes. Super pause sequences (**Fig. 4A**) could serve to tightly regulate the timing and competition between premature termination and P-TEFb dependent activation. If DSIF or NELF fail to associate with RNA polymerase II at super pause sites, RNA polymerase II may be more prone to enter the sidetracked state. This could favor premature termination/attenuation (**Fig. 7D**). Whether DSIF-NELF favor or disfavor the sidetracked state remains to be explored. The interplay between sequence and factor control at pause sites could support the rate of release into transcription elongation or premature termination.

In summary, GATO-seq is a highly sensitive and modular technique that can allow for the study of all aspects of transcriptional regulation across species and will serve as a complement to cell-based approaches. We envision that GATO-seq will serve as a platform for bottom-up reconstitution of transcriptional processes. The GATO-seq approach, together with cell-based, biochemical, and biophysical assays, will enable a more complete and mechanistic understanding of gene regulation.

### Limitations of the study

Topological constraints are known to affect transcription elongation^98–103^. The experiments shown here were performed on linear DNA substrates lacking chromatin and do not account for these effects. Our experiments use a viral promoter, and thus the effect of native promoter sequence is not examined here. We additionally are missing initiation factors such as TFIID and Mediator, which may affect pause duration or pause site selection^37,50,63^. Only three time points were collected in our GATO-seq experiments, and finer time sampling will help in better modelling pause behaviors. The use of three time points for determining pausing half-life can result in under-or over-estimation of pausing half-life (**Fig. S2B**). Including five time points results in more robust pause half-life estimation. Additionally, our assay does not account for premature termination, and future assays could address this by including factors like Integrator or ARMC5. As observed in previously published gel-based assays, only a fraction of elongation complexes produced full-length transcripts (maximally 50%) ^59,104–106^. This could indicate the presence of prematurely terminated products or strongly paused elongation complexes. Such incomplete transcripts could be additionally generated by incomplete DNA templates, RNA shredding during library preparation or pyrophosphorolysis. Our method cannot distinguish between these products. Finally, we leverage the use of poly-A tails to facilitate library preparation. This could result in depletion of pause sites in which the terminal nucleotide was an adenosine. A combination of other poly-nucleotide tails, or direct 3’end ligation to preadenylated adapters could circumvent this limitation.

## Acknowledgments

We thank all past and present members of the Vos lab for support, discussion, and comments on the manuscript. Cryo-EM specimens were prepared, and data was collected at the Cryo-EM Facility at MIT.nano. We thank S. Sterling, J. Podgorski and N. Swanson for support at MIT.nano, L. Farnung for sharing reagents and critical discussions, E. Guzman-Cerezo for initial code preparation, L. Sowin for initial protein purifications and cloning assistance, K. Adelman and C. Mimoso for sharing HEK293T PRO-seq analysis, and C. Burge and H. Kurka Margolis for critical discussions.

## Funding

National Institutes of Health grant DP2-GM146254 (SMV)

Freeman Hrabowski Scholar of the Howard Hughes Medical Institute (SMV)

Swiss National Science Foundation, Early Postdoc Mobility Fellowship (RVN)

## Author contributions

Conceptualization: RVN, SMV

Methodology: RVN, SMV

Investigation: RVN, SK, SMV

Visualization: RVN, SMV

Funding acquisition: SMV

Project administration: SMV

Supervision: SMV

Writing – original draft: RVN, SMV

Writing – review & editing: RVN, SMV

## Declaration of interests

Authors declare that they have no competing interests. Supplementary Information is available for this paper.

### Lead contact

Correspondence and requests for materials should be addressed to.

## Materials availability

Materials available upon request from Seychelle M. Vos.

## Data and code availability

Structure models coordinates are deposited in the PDB (XXXX sidetracked and YYYY pretranslocated) and the cryo-EM maps are deposited into the Electron Microscopy Data Bank (EMDB): EMD-XXXX and EMD-ZZZZ. Original code is available at Zenodo repository with the following link XXX Oxford nanopore direct RNA sequencing reads are available at SRA under the accession number PRJNAXXXX. Processed GATO-seq signal data is available at the GEO under the accession number GSEXXXX.

## Methods

### Experimental model and subject details

Bacterial strains. *E. coli* strains DH5α (New England Biolabs), BL21 (DE3) RIL (Merck), DH10EMBacY (Geneva Biotech) were grown in standard LB or 2xYT media at 37 ºC supplemented with ampicillin (100 µg/mL), gentamycin (10 µg/mL), kanamycin (30 µg/mL), or chloramphenicol (34 µg/mL).

Insect cells Sf9 (Expression Systems), Sf21 (Expression Systems), and *Trichoplusia ni* Hi5 (Expression Systems) cells were cultured in ESF 921 media (Expression Systems) at 27ºC.

Porcine (*Sus scrofa*) thymus tissue was obtained from Pel-Freez Biologicals. The tissues were harvested from young and healthy pigs, frozen at -20ºC, and stored at -80ºC prior to protein purification.

### Cloning and protein expression

DSIF was cloned and expressed as previously described ^108^. Briefly, *E. coli* BL21(DE3) RIL transformed with the DSIF expression plasmid were grown at 37ºC. Protein expression was induced when cells reached an OD 600 nm of 0.5 by adding 0.5 mM IPTG. After 3 hours of expression at 37ºC, cells were collected by centrifugation and resuspended in DSIF lysis buffer (50 mM Na-HEPES pH 7.4, 500 mM NaCl, 30 mM Imidazole pH 8, 10% (v/v) glycerol, 5 mM β-mercaptoethanol, 0.284 µg/mL leupeptin, 1.37 µg/mL pepstatin A, 0.17 mg/mL PMSF, and 0.33 mg/mL benzamidine), flash frozen in liquid nitrogen, and stored at -80ºC until protein purification.

NELF was cloned as described ^109^. Hi5 cells were infected with baculovirus carrying NELF-A, -B, His6-TEV-D and -E subunits, and protein expression was carried out for 48 hours after infection. Cells were collected by centrifugation and resuspended in lysis buffer (300 mM NaCl, 20 mM Na-HEPES pH 7.4, 10% (v/v) glycerol, 30 mM imidazole pH 8.0, 5 mM β-mercaptoethanol, 0.284 mg/mL leupeptin, 1.37 mg/mL pepstatin A, 0.17 mg/mL PMSF, and 0.33 mg/mL benzamidine), flash frozen in liquid nitrogen, and stored at -80 ºC until protein purification.

TFIIS and TFIIS DE282-283AA were cloned and expressed as previously described ^54^. *E. coli* BL21(DE3) RIL transformed with the TFIIS or TFIIS DE282-283AA expression plasmid were grown at 37ºC. Protein expression was induced when cells reached an OD 600 nm of 0.5 by adding 0.5 mM IPTG. Cells were grown for an additional 16h at 18^°^C, collected by centrifugation, resuspended in TFIIS lysis buffer (800 mM NaCl, 20 mM Tris-HCl pH 7.9, 10% (v/v) glycerol, 30 mM imidazole pH 8.0, 1 mM DTT, 0.284 µg/mL leupeptin, 1.37 µg/mL pepstatin A, 0.17 mg/mL PMSF and 0.33 mg/mL benzamidine), flash frozen in liquid nitrogen, and stored at -80ºC until protein purification.

Human elongation factors, SPT6, PAF, RTF1 and P-TEFb were cloned, expressed and purified as previously described in (*43*).

Human transcription initiation factors were cloned in full-length form unless otherwise noted. The following cloning and expression procedures were followed for initiation factors that were expressed using baculovirus: TFIIA subunit 1 was inserted in 438-A (AddGene: 55218) with a C-terminal TEV cleavable 6X histidine tag. TFIIA subunit 2 was cloned in 438-A. Both genes were combined into a single MacroBac vector to produce a final expression vector ^110^. TBP was cloned into plasmid 438-B (AddGene: 55219) containing a TEV cleavable 6x histidine N-terminal tag. TFIIH was assembled in three vectors. The first vector contained subunits GTF2H1, 2, 3, 4, 5 and ERCC3. GTF2H1, 2, 4 and ERCC3 bore N-terminal 6x histidine tags followed by a TEV cleavage site (438-B). GTF2H3 and GTF2H5 were cloned in 438-A. The 6 subunits were combined into pBIG2ab ^111^. ERCC2 was cloned in vector 438-C resulting in a N-terminal 6x Histidine-maltose binding protein (MBP) tag followed by a TEV cleavage site (AddGene: 55220). The CDK activating kinase (CAK) module subunits CCNH, CDK7 and MAT1 were individually cloned into vectors 438-C, B and A, respectively. The three subunits were combined into the pBig1a vector (AddGene: 80611). These baculovirus constructs were transformed into *E. coli* DH10EMBacY for bacmid generation. Positive colonies containing the bacmid were isolated and amplified. Bacmids were isolated by alkaline lysis and ethanol precipitation and transfected into adherent Sf9 cells (Expression Systems) to produce V0 virus. V0 virus was amplified into V1virus in Sf9 or Sf21 cells. For large scale expression, V1 virus was used to infect Hi5 cells. Typically, 600 mL of Hi5 cells at density of 1 million cells/mL were infected. After 72 hours of protein expression, cells were harvested by centrifugation and resuspended in GTF lysis buffer (25 mM Na-HEPES pH 7.4, 500 mM KCl, 30 mM Imidazole pH 8.0, 10% v/v glycerol, 5 mM β-mercaptoethanol, 0.284 mg/mL leupeptin, 1.37 mg/mL pepstatin A, 0.17 mg/mL PMSF, and 0.33 mg/mL benzamidine). TFIIH was resuspended in TFIIH lysis buffer (400 mM KCl, 20 mM Na-HEPES pH 7.0, 20% (v/v) glycerol, 30 mM imidazole pH 8.0, 5 mM β-mercaptoethanol, 0.284 mg/mL leupeptin, 1.37 mg/mL pepstatin A, 0.17 mg/mL PMSF, and 0.33 mg/mL benzamidine). Resuspended cells were snap frozen in liquid nitrogen and stored at -80ºC prior to purification.

The following cloning and expression procedures were followed for human transcription initiation factors that were expressed in *E. coli*. TFIIB was cloned into a modified pET28b vector behind a N-terminal 6x His followed by a TEV cleavage site (AddGene: 69865-3). TFIIE subunit 1 was cloned in the untagged 14-A vector (AddGene: 48307). TFIIE subunit 2 was cloned in vector 14-B behind a N-terminal TEV cleavable 6x His tag (AddGene: 48308). Vectors were combined to produce a single vector for expression. TFIIF subunit 1 was inserted in vector 14-B and subunit 2 in vector 14-A, and the subunits were combined into a single vector.

TFIIF and TFIIE were expressed in *E. coli* BL21(DE3) RIL. Cells were grown at 37ºC until reaching an OD 600 nm of 0.5. Protein expression was induced by adding 0.5 mM IPTG, and cells were grown for an additional 16 hrs at 18^°^C. TFIIB was expressed in *E. coli* LOBSTR-BL21(DE3)-RIL cells (Kerafast) grown at 37 ^°^C in auto-induction media. The temperature was decreased to 18ºC when cells reached an OD 600 nm of 0.4 and were grown for an additional 12h. After expression, cells were collected by centrifugation and resuspended in GTF lysis buffer. All the resuspended cells were flash frozen in liquid nitrogen and stored at -80^°^C until protein purification.

### Protein purification

*S. scrofa* RNA polymerase II was purified from thymus tissue as previously described ^54,55,112,113^. Briefly, 500 g of frozen thymus was pulverized by hammering and homogenized by blending in 2 L RNA polymerase II lysis buffer (20 mM Tris-HCl pH 7.9 at 4^°^C, 10% v/v glycerol, 1 mM EDTA, 25 µM ZnCl_2_, 0.284 µg/mL leupeptin, 1.37 µg/mL pepstatin A, 0.17 mg/mL PMSF, and 0.33 mg/mL benzamidine). Homogenate was clarified by centrifugation and precipitated with 0.04% v/v polyethylenimine (Sigma). The precipitated material was washed once with RNA polymerase II lysis buffer and protein was extracted from the pellets by resuspending in RNA polymerase II lysis buffer containing 150 mM ammonium sulfate (Sigma) followed by centrifugation. The supernatant was applied to a 250 mL MacroQ column (Bio-Rad) equilibrated in RNA polymerase II lysis buffer containing 200 mM ammonium sulfate. The column was washed, and protein was eluted in RNA polymerase II lysis buffer containing 400 mM ammonium sulfate. Peak fractions were pooled and precipitated by adding ammonium sulfate to 50% saturation at 4^°^C. Precipitant was collected by centrifugation and resuspended in RNA polymerase II Lysis buffer containing 1 mM DTT and adjusted to a final concentration of 150 mM ammonium sulfate. The sample was applied to a 5 mL gravity column packed with 8WG16 anti-RNA polymerase II CTD antibody (Sigma-Millipore) conjugated to agarose resin, washed with RNA polymerase II lysis buffer containing 500 mM Ammonium sulfate and eluted with the same buffer supplemented with 50% v/v glycerol at room temperature. Elution fractions were analyzed by SDS-PAGE and RNA polymerase II was further purified by applying the fractions to a 1 mL UnoQ ion exchange column (BioRad) equilibrated with RNA polymerase II lysis buffer containing 2 mM DTT and 100 mM Ammonium sulfate. Protein was eluted via a gradient from 100 to 500 mM ammonium sulfate in RNA polymerase II lysis buffer. Peak fractions were analyzed by SDS-PAGE. Peak fractions were concentrated using a 100 kDa MWCO Amicon Ultra Centrifugal Filter (Sigma-Aldrich), dialyzed in RNA polymerase II storage buffer (25 mM Na-HEPES pH 7.4, 150 mM NaCl and 10 mM DTT), aliquoted, flash frozen in liquid nitrogen, and stored at -80^°^C.

WT TFIIS and TFIIS DE282-283AA were purified from 6L of culture ^54^. Cells resuspended in TFIIS lysis buffer were lysed by sonication. The lysate was clarified by centrifugation. The supernatant was applied to a 5 mL HisTrap column (Cytiva) equilibrated in lysis buffer. The column was washed with lysis buffer and equilibrated in low salt buffer (400 mM NaCl, 20 mM Tris-HCl pH 7.9, 10% (v/v) glycerol, 30 mM imidazole pH 8.0, 1 mM DTT, 0.284 µg/mL leupeptin, 1.37 µg/mL pepstatin A, 0.17 mg/mL PMSF, and 0.33 mg/mL benzamidine) until no additional absorbance was detected at 280 nm. The protein was eluted from the nickel column by a gradient over 6 column volumes with low salt buffer supplemented with 500 mM imidazole pH 8.0. Peak fractions were analyzed by SDS-PAGE and Coomassie staining. Fractions containing TFIIS were pooled and mixed with 1.5 mg 6xHis tagged TEV protease and dialyzed overnight against low salt buffer in SnakeSkin dialysis tubing (10 kDa MWCO). The protein was removed from dialysis tubing and uncleaved protein and TEV protease were removed by applying the sample to a 5 mL HisTrap column equilibrated in low salt buffer. The flow through was collected, concentrated in a 10 kDa MWCO Amicon Ultra Centrifugal Filter (Sigma-Aldrich), and applied to a HiLoad S75 16/1600 pg column (Cytiva) equilibrated in 400 mM NaCl, 20 mM Tris-HCl pH 7.9, 10% (v/v) glycerol, and 1 mM DTT. Protein purity was determined by SDS-PAGE. Peak fractions were concentrated in 10 kDa MWCO Amicon Ultra Centrifugal Filter (Signa-Aldrich). The protein was stored in a buffer containing 400 mM NaCl, 20 mM Tris-HCl pH 7.9, 30% (v/v) glycerol, and 1 mM DTT. The protein was aliquoted, flash-frozen in liquid nitrogen, and stored at -80°C.

DSIF was purified from 6 L of *E. coli* BL21 (DE3) RIL ^108^. Resuspended cells were lysed by sonication and clarified by centrifugation. The clarified lysate was filtered through a 0.45 µm syringe filter and applied to a 5 mL HisTrap HP column (Cytiva) equilibrated in lysis buffer (500 mM NaCl, 50 mM Na-HEPES pH 7.4, 10% (v/v) glycerol, 50 mM imidazole pH 8.0, 1 mM DTT, 0.284 µg/mL leupeptin, 1.37 µg/mL pepstatin A, 0.17 mg/mL PMSF, and 0.33 mg/mL benzamidine). The column was washed with 10 column volumes of lysis buffer, followed by 3 column volumes of lysis buffer supplemented with 1000 mM NaCl. Another wash step with 3 column volumes of lysis buffer was performed before eluting the protein over a gradient using lysis buffer containing 500 mM imidazole pH 8.0. Peak fractions containing DSIF were pooled, mixed with 3C protease with an uncleavable 6×His tag, and dialyzed overnight in 10 kDa MWCO SnakeSkin dialysis tubing (Thermo Scientific) against Q buffer (300 mM NaCl, 50 mM Na-HEPES pH 7.4, 10% (v/v) glycerol, 30 mM imidazole pH 8.0, and 1 mM DTT). The protein was collected and applied to a 5 mL HisTrap and a 5 mL HiTrap Q column (Cytiva) connected in tandem and equilibrated in Q buffer to remove the 3C protease, the His tag, uncleaved protein, and protein lacking the acidic N-terminal region of SPT5. The tandem column was washed with 5 column volumes of Q buffer, after which the HisTrap column was removed. The HiTrap Q column was eluted over a gradient using Q buffer containing 1000 mM NaCl. Peak fractions were pooled, and protein purity was assessed by SDS-PAGE followed by Coomassie staining. Pure DSIF was concentrated using 50 kDa MWCO Amicon Ultra Centrifugal Filters (Sigma-Aldrich) and applied to a HiLoad S200 16/600 pg column (Cytiva) equilibrated in 500 mM NaCl, 20 mM Na-HEPES pH 7.4, 10% (v/v) glycerol, and 1 mM DTT. Protein purity was assessed by SDS-PAGE and Coomassie staining. Pure fractions containing full-length DSIF were concentrated using 50 kDa MWCO Amicon Ultra Centrifugal Filters (Sigma-Aldrich). The protein concentration was determined by measuring absorbance at 280 nm and using the predicted extinction coefficient for the complex. Purified DSIF was aliquoted, flash-frozen, and stored at -80 °C.

NELF was purified as previously described ^109^. Hi5 cells (1.2 L) were harvested and lysed by sonication. The lysate was clarified by centrifugation and filtered through 0.45 µm syringe filters. The clarified lysate was applied to a 5 mL HisTrap HP column (Cytiva) equilibrated in lysis buffer (300 mM NaCl, 20 mM Na-HEPES pH 7.4, 10% (v/v) glycerol, 30 mM imidazole pH 8.0, 1 mM DTT, 0.284 µg/mL leupeptin, 1.37 µg/mL pepstatin A, 0.17 mg/mL PMSF, and 0.33 mg/mL benzamidine). The column was washed with 10 column volumes of lysis buffer, followed by 3 column volumes of lysis buffer supplemented with 800 mM NaCl. Another wash step with 3 column volumes of low-salt buffer (150 mM NaCl, 20 mM Na-HEPES pH 7.4, 10% (v/v) glycerol, 50 mM imidazole pH 8.0, 1 mM DTT, 0.284 µg/mL leupeptin, 1.37 µg/mL pepstatin A, 0.17 mg/mL PMSF, and 0.33 mg/mL benzamidine) was performed. A 5 mL HiTrap Q column (Cytiva) equilibrated in low-salt buffer was then connected in series, and the protein was eluted over a gradient using low-salt buffer supplemented with 500 mM imidazole. The columns were washed with low-salt buffer before detaching the HisTrap column. NELF was subsequently eluted from the HiTrap Q column using a gradient to 800 mM NaCl in a buffer containing 20 mM NaHEPES pH 7.4, 10% (v/v) glycerol, 1 mM DTT. Peak fractions containing NELF were pooled, mixed with 1.5 mg TEV protease, and dialyzed overnight in 10 kDa MWCO SnakeSkin dialysis tubing (Thermo Scientific) against lysis buffer. The protein was collected and applied to a 5 mL HisTrap column to remove TEV protease, uncleaved protein, and the His tag. The flowthrough was concentrated using 50 kDa MWCO Amicon Ultra Centrifugal Filters (Sigma-Aldrich). The concentrated sample was applied to a HiLoad S200 16/600 pg column (Cytiva) equilibrated in 150 mM NaCl, 20 mM Na-HEPES pH 7.4, 10% (v/v) glycerol, and 1 mM DTT. Protein integrity and purity were assessed by SDS-PAGE followed by Coomassie staining. Pure fractions were concentrated using 50 kDa MWCO Amicon Ultra Centrifugal Filters (Sigma-Aldrich). The protein concentration was determined by measuring absorbance at 280 nm and the predicted extinction coefficient for the complex. Purified NELF was aliquoted, flash-frozen, and stored at -80 °C.

TFIIA was purified from 1.6 L of Hi5 cells ^114^. Cells were lysed by sonication. The lysate was clarified by centrifugation and filtered through 0.45 µm syringe filters. The clarified lysate was applied to a 5 mL HisTrap HP column (Cytiva) equilibrated in lysis buffer. The column was washed with 10 column volumes (CV) of lysis buffer, followed by 5 CV of lysis buffer containing 1000 mM KCl, and 5 CV of lysis buffer containing 100 mM KCl. A 5 mL HiTrap Heparin HP column (Cytiva) in series with a 5 mL HiTrap Q HP column (Cytiva) was attached to the base of the HisTrap column. TFIIA was eluted from the HisTrap column using a gradient up to 500 mM imidazole. The HisTrap column was then detached, and the tandem Heparin-Q column was washed with 10 CV of buffer containing 100 mM KCl, 25 mM Na-HEPES pH 7.4, 10% (v/v) glycerol, and 1 mM DTT. The Heparin column was subsequently removed, and TFIIA was eluted from the Q column with a step gradient, first using a buffer containing 150 mM KCl, followed by a gradient to 750 mM KCl. Peak fractions were analyzed by SDS-PAGE, and those containing TFIIA were pooled and concentrated using a 10 kDa MWCO Amicon Ultra Centrifugal Filter (Sigma-Aldrich). The sample was then applied to a HiLoad Superdex 200 pg 16/600 column (Cytiva) equilibrated in 100 mM KCl, 25 mM Na-HEPES pH 7.4, 10% (v/v) glycerol, and 1 mM DTT. The final peak fractions were analyzed by SDS-PAGE, pooled, and concentrated using a 10 kDa MWCO Amicon Ultra Centrifugal Filter (Sigma-Aldrich). Protein concentration was determined by measuring absorbance at 280 nm and the theoretical extinction coefficient. Purified TFIIA was aliquoted, flash-frozen, and stored at -80 °C.

TFIIB was purified from 6 L of expression ^114^. After harvesting, cells were resuspended in GTF lysis buffer, sonicated, and centrifuged. The clarified lysate was applied to a 5 mL HisTrap column equilibrated in lysis buffer. The column was washed with 10 column volumes (CV) of lysis buffer, followed by a high salt wash with 5 CV of GTF lysis buffer supplemented with 1000 mM KCl. The column was then washed with 5 CV of lysis buffer. The protein was eluted with a gradient of imidazole to 500 mM. Peak fractions were analyzed by SDS-PAGE. Fractions containing TFIIB were pooled, mixed with 1.5 mg of TEV protease, and dialyzed overnight against dialysis buffer containing 200 mM KCl, 25 mM Na-HEPES pH 7.4, 10% (v/v) glycerol, and 1 mM DTT. TEV protease and uncleaved protein were separated by flowing the dialyzed sample over a 5 mL HisTrap column equilibrated in lysis buffer. The flowthrough containing cleaved TFIIB was applied to a 5 mL HiTrap SP (Cytiva) column equilibrated in lysis buffer. After washing with 10 CV of lysis buffer, TFIIB was eluted by developing the column over a KCl gradient to 1000 mM. Peak fractions were analyzed by SDS-PAGE, and the fractions containing TFIIB were concentrated using a 10 kDa MWCO Amicon Ultra Centrifugal Filter (Sigma-Aldrich). The concentrated protein was applied to a HiLoad Superdex 75 pg 16/600 column equilibrated in storage buffer (200 mM KCl, 25 mM Na-HEPES pH 7.4, 10% (v/v) glycerol, and 0.5 mM TCEP). The peak fractions were visualized by SDS-PAGE, and then concentrated using a 10 kDa MWCO Amicon Ultra Centrifugal Filter (Sigma-Aldrich). The final concentration was determined by measuring the absorbance at 280 nm and the theoretical extinction coefficient. TFIIB was then aliquoted, flash frozen in liquid nitrogen, and stored at - 80°C.

Pelleted and resuspended Hi5 cells (600 mL expression) expressing His-tagged TBP were lysed by sonication ^114^. The lysate was clarified by centrifugation and filtration through 0.45 µm syringe filters. The clarified lysate was applied to a 5 mL HisTrap column equilibrated in GTF lysis buffer. The column was washed with 10 column volumes (CV) of GTF lysis buffer, and TBP was eluted using an imidazole gradient to 500 mM in GTF lysis buffer. The peak fractions were pooled after SDS-PAGE, mixed with1.5 mg of TEV protease, and dialyzed overnight against dialysis buffer containing 150 mM KCl, 25 mM Na-HEPES (pH 7.5), 10% (v/v) glycerol, and 1 mM DTT. The sample was collected and applied to a HisTrap column equilibrated in dialysis buffer to remove uncleaved protein, TEV protease, and the free His6 tag. The flowthrough was collected and applied to a HiTrap SP column equilibrated in dialysis buffer. After washing the column with 10 CV of dialysis buffer, the protein was eluted with a KCl gradient to 1000 mM. The peak fractions were pooled and concentrated using a 10 kDa MWCO Amicon Ultra Centrifugal Filter (Sigma-Aldrich). TBP was then applied to a Superdex 200 GL 10/300 column equilibrated in storage buffer (200 mM KCl, 25 mM Na-HEPES pH 7.5, 10% (v/v) glycerol, and 0.5 mM TCEP). The fractions containing TBP were analyzed by SDS-PAGE and subsequently concentrated using a 10 kDa MWCO Amicon Ultra Centrifugal Filter (Sigma-Aldrich). Protein concentration was estimated using the extinction coefficient and absorbance at 280 nm. TBP was aliquoted, flash frozen in liquid nitrogen, and stored at -80°C.

Resuspended *E. coli* cells expressing TFIIF (6 L) were lysed by sonication^114^. The lysate was clarified by centrifugation and filtration through 45 µm syringe filters. The clarified lysate was applied to a 5 mL HisTrap column equilibrated in GTF lysis buffer. After washing the column with 10 column volumes (CV) of lysis buffer, the sample was further washed with GTF lysis buffer supplemented with 1000 mM KCl, followed by GTF lysis buffer. Protein was eluted using an imidazole gradient up to 500 mM. The peak fractions were analyzed by SDS-PAGE and fractions containing TFIIF were pooled. The pooled protein was mixed with 1.5 mg of TEV protease and dialyzed overnight against dialysis buffer (20 mM Na-HEPES pH 7.5, 100 mM KCl, 10% (v/v) glycerol, 5 µM ZnCl_2_, and 1 mM DTT). The cleaved protein was collected and applied to a HisTrap column equilibrated in dialysis buffer. The flowthrough was collected and applied to a HiTrap SP column equilibrated in dialysis buffer. The column was washed with 10 CV of dialysis buffer, and the protein was eluted using a KCl gradient to 1000 mM. The peak fractions were analyzed by SDS-PAGE, concentrated using a 30 kDa MWCO Amicon Ultra Centrifugal Filter (Sigma-Aldrich), and applied to a HiLoad Superdex 200 pg column equilibrated in storage buffer (20 mM Na-HEPES pH 7.5, 200 mM KCl, 10% (v/v) glycerol, and 0.5 mM TCEP). The eluted protein was visualized by SDS-PAGE and further concentrated using a 30 kDa MWCO Amicon Ultra Centrifugal Filter (Sigma-Aldrich). TFIIF was aliquoted, flash frozen in liquid nitrogen, and stored at -80°C. Protein concentration was determined by measuring the absorbance at 280 nm and the theoretical extinction coefficient.

Resuspended *E. coli* cells expressing TFIIE (6L) were lysed by sonication ^114^, and the lysate was clarified by centrifugation and by syringe filtration using a 0.45 µm syringe filter. The clarified lysate was applied to a 5 mL HisTrap column equilibrated in lysis buffer. The column was washed with 10 CV of GTF lysis buffer, followed by 5 CV of lysis buffer supplemented with 1000 mM NaCl, and lysis buffer. The protein was eluted using an imidazole gradient to 500 mM. The peak fractions were analyzed by SDS-PAGE, pooled and mixed with 1.5 mg TEV protease, and dialyzed overnight against dialysis buffer (200 mM NaCl, 50 mM Tris-HCl pH 8.0, 10% (v/v) glycerol, 50 mM imidazole pH 8.0, and 1 mM DTT). After dialysis, the sample was collected and applied to a 5 mL HisTrap column equilibrated in dialysis buffer. The flowthrough was collected and loaded onto a 5 mL Heparin column equilibrated in dialysis buffer. The column was washed with 10 CV of dialysis buffer supplemented with 250 mM NaCl, and the protein was eluted using a NaCl gradient to 1500 mM. The peak fractions were analyzed by SDS-PAGE, pooled, and concentrated using a 10 kDa MWCO Amicon Ultra Centrifugal Filter (Sigma-Aldrich). TFIIE was further purified on a HiLoad Superdex 200 pg 16/600 column equilibrated in storage buffer (200 mM NaCl, 25 mM Na-HEPES pH 7.5, 10% (v/v) glycerol, 50 mM imidazole pH 8.0, and 0.5 mM TCEP). The peak fractions were analyzed by SDS-PAGE, pooled, and concentrated using a 30 kDa MWCO Amicon Ultra Centrifugal Filter (Sigma-Aldrich). Protein concentration was determined by measuring the absorbance at 280 nm and the theoretical extinction coefficient. TFIIE was aliquoted, flash frozen in liquid nitrogen, and stored at -80°C. Resuspended Hi5 cells expressing TFIIH were lysed by sonication, the lysate was clarified by centrifugation and filtration through a syringe filter (0.45 µm) ^115^. The clarified lysate was applied to a 5 mL HisTrap column equilibrated in TFIIH lysis buffer. The column was washed with 10 column volumes (CV) of TFIIH lysis buffer, followed by a wash with TFIIH lysis buffer supplemented with 800 mM KCl, and 5 CV of TFIIH lysis buffer. The protein was eluted using an imidazole gradient to 500 mM onto a 15 mL amylose column connected in series. After disconnecting the HisTrap column, the amylose column was washed with 5 CV of amylose wash buffer (20 mM Na-HEPES pH 7.0, 400 mM KCl, 20% (v/v) glycerol, and 1 mM DTT). TFIIH was eluted using a step gradient with amylose wash buffer containing 100 mM maltose. The peak fractions were analyzed by SDSPAGE and pooled. The pooled fraction was diluted with TFIIH dilution buffer (20 mM Na-HEPES pH 7.0, 20% (v/v) glycerol, and 1 mM DTT) to adjust the KCl concentration to 250 mM. The sample was then applied to a heparin column equilibrated in heparin wash buffer (20 mM Na-HEPES pH 7.0, 250 mM KCl, 20% (v/v) glycerol, and 1 mM DTT). The column was washed with 10 CV of heparin wash buffer, and TFIIH was eluted using a KCl gradient to 800 mM. The peak fractions containing stoichiometric amounts of TFIIH core subunits were applied to a Superose 6 GL 10/300 column equilibrated in TFIIH storage buffer (20 mM Na-HEPES pH 7.0, 400 mM KCl, 20% (v/v) glycerol, and 1 mM TCEP). The peak fractions were analyzed by SDS-PAGE, pooled, and concentrated using a 100 kDa MWCO Amicon Ultra (Sigma-Aldrich). Protein concentration was determined by measuring the absorbance at 280 nm and the theoretical extinction coefficient of the combined subunits. The concentrated TFIIH was aliquoted, flash frozen in liquid nitrogen, and stored at -80°C.

Hi5 cells expressing CAK were harvested by centrifugation and resuspended in CAK lysis buffer (400 mM KCl, 25 mM Na-HEPES pH 7.4, 10% (v/v) glycerol, 30 mM imidazole pH 8.0, 1 mM DTT, 0.284 µg/mL leupeptin, 1.37 µg/mL pepstatin A, 0.17 mg/mL PMSF, and 0.33 mg/mL benzamidine)^115^. Cells were lysed by sonication, and the lysate was clarified by centrifugation and filtration through a 0.45 µm syringe filter. The clarified lysate was applied to a 5 mL HisTrap column equilibrated in CAK lysis buffer. The column was washed with 10 column volumes (CV) of CAK lysis buffer, and protein was eluted using an imidazole gradient to 500 mM onto a 15 mL amylose column connected in series. After disconnecting the HisTrap column, the amylose column was washed with 10 CV of amylose wash buffer (400 mM KCl, 25 mM Na-HEPES pH 7.4, 10% (v/v) glycerol, and 1 mM DTT) and eluted using a step gradient with amylose wash buffer containing 100 mM maltose. The peak fractions were analyzed by SDS-PAGE and concentrated using a 30 kDa MWCO Amicon Ultra Centrifugal Filter (Sigma-Aldrich). The sample was applied to a HiLoad Superdex 200 pg 16/600 column equilibrated in CAK storage buffer (400 mM KCl, 25 mM Na-HEPES pH 7.4, 10% (v/v) glycerol, and 0.5 mM TCEP). The peak fractions were analyzed by SDS-PAGE, and those containing stoichiometric amounts of CAK subunits were pooled and concentrated in a 30 kDa MWCO Amicon Ultra Centrifugal Filter (Sigma-Aldrich). Protein concentration was determined by measuring the absorbance at 280 nm and the theoretical extinction coefficient of the combined subunits. The purified protein was aliquoted, flash frozen in liquid nitrogen, and stored at -80°C.

### RNA extension assays

All oligos were purchased from Sigma-Aldrich, resuspended in water (100 µM), flash-frozen in liquid nitrogen, and stored at -80°C. RNA extension assays were performed with perfectly complementary scaffolds. Super pause consensus sequences were: DNA template 5′-CCA CTG GAA GAT CTG AAT TTG CGG CAG CAG CTC CGC CGG ACG TAC TGA CC -3′, DNA non-template 5′-GGT CAG TAC GTC CGG CGG AGC TGC TGC CGC AAA TTC AGA TCT TCC AGT GG -3′, RNA 5′-6-FAM-UUU UUU GGC GGA GCU-3′. Mutated Super pause sequences were: DNA template 5′-CCA CTG GAA GAT CTG AAT TTG CGG AAG CAG CTC CGC CGG ACG TAC TGA CC-3′, DNA non-template 5′-GGT CAG TAC GTC CGG CGG AGC TGC TTC CGC AAA TTC AGA TCT TCC AGT GG-3′, RNA 5′-6-FAM-UUU UUU GGC GGA GCU -3′. The 9-base-pair DNA-RNA hybrid is preceded by 13 nucleotides of upstream DNA and succeeded by 28 nucleotides of downstream DNA. RNA and template DNA were mixed in equimolar ratios and were annealed by incubating the nucleic acids at 95°C for 5 min and then decreasing the temperature by 1°C min^−1^ steps to a final temperature of 30°C in a thermocycler in a buffer containing 100 mM NaCl, 20 mM Na-HEPES pH 7.4, 3 mM MgCl_2_, and 10% (v/v) glycerol. All concentrations correspond to the final concentrations used in the assay. *S. scrofa* RNA polymerase II (100 nM) and the RNA-DNA template hybrid (100 nM) were incubated for 10 min at 30 °C, shaking at 300 rpm. The non-template DNA (100 nM) was added, and the reactions were incubated for another 10 min. Factors were diluted in protein dilution buffer (300 mM NaCl, 20 mM Na-HEPES pH 7.4, 10% (v/v) glycerol and 1 mM DTT) and added to RNA polymerase II elongation complexes at a concentration of 300 nM. The reactions were then diluted to achieve final assay conditions of 100 mM NaCl, 20 mM Na-HEPES pH 7.4, 3 mM MgCl_2_, 4% (v/v) glycerol, 1 mM DTT, 25 µM ZnCl_2_ and further incubated for 10 min at 30 °C. Transcription reactions were initiated by adding NTPs at a final concentration of 10 µM or 1 mM to permit elongation. Reactions (5 µL) were quenched after 0-10 min in 5 µL 2x Stop buffer (6.4 M urea, 50 mM EDTA pH 8.0, 1x TBE buffer and 4 µg of proteinase K (New England Biolabs)). Quenched samples were incubated for 30 min at 37°C and were separated by denaturing gel electrophoresis (5 µL of sample applied to an 8 M urea, 1x TBE, 20% Bis-Tris acrylamide 19:1 gel run in 0.2x TBE buffer at 300V for 160 min). Products were visualized using the 6-FAM label on a Typhoon 9500 FLA Imager (GE Healthcare Life Sciences) at 900 PMT. Gel images were quantified using ImageJ (v1.53K). A 0.48 × 1.98 cm box surrounding each lane was created to obtain lane plots. A baseline was manually placed, and the tracing tool was used to obtain the total integrated density for each band. The integrated density of the +3 pause band was normalized relative to the intensity of the starting RNA band, by dividing the pause band integrated density value by that of the starting RNA band. Graphs were prepared in GraphPad Prism 10. Each point represents the mean intensity from three individual replicates. Line graph error bars reflect standard deviation of replicates.

### Sequences retrieving for GATO-seq library templates design

The top 1,000 sequences from human hg38.p14 genome assemble, ranked by TSS focus score were selected using GetGenAnnotation pipeline (https://zenodo.org/records/5519928) from WT 293T PRO-seq and RNA-seq data ^116^, kindly provided by Claudia Mimoso and Karen Adelman (Harvard Medical School). Genome coordinates were used to retrieve sequences comprising the regions from TSS to gene position +261 from the genome data bank using a custom Python script available at Zenodo

### AdML plasmid library preparation

Retrieved sequences were flanked with a 20 bp 5′ cloning adaptor: 5′-GAA CCC ACT CGG AGC CAG CA -3′ and a 3′ cloning adaptor: 5′-GCG GTG GAG CTC CAA TTC G -3′ and synthesized as an oligo pool by silicon-based DNA synthesis platform (Twist Bioscience). The oligo pool was amplified following the manufacturer’s instructions with the following DNA primers: forward 5′-CAC TCC ACT CGG AGC CAG CA-3′ and reverse 5′-GCG GTG GAG CTC CAA TTC G -3′. The size distribution of amplified DNA fragments was estimated by fragment analysis. The fragment pool (300 ng) was cloned by Gibson assembly into 1 µg of linearized pBlueScript vector containing the AdML promoter and 5’ adaptor sequence. Cloned vectors were transformed into 500 µL NEB® 5-alpha Competent *E. coli* (High Efficiency) (New England Biolabs) in 10 separate reactions following the manufacturer’s instructions and plated on 100 LB agar plates supplemented with carbenicillin. Plates were incubated overnight at 37 ^°^C. Colonies were collected and grown for 8 h at 37 ^°^C in 20 L LB media split in 20 flasks supplemented with ampicillin, shaking at 160 rpm. Biomass was collected by centrifugation and cells were opened by alkaline lysis. The plasmid library was extracted by phenol-chloroform-isoamyl alcohol and precipitated with 2 volumes of 100% ethanol. The plasmid pellets were resuspended in 10 mL nuclease-free H_2_O and applied in 3.3 mL volume injections onto a HiPrep 16/600 Sephacryl S-500 size exclusion column (Cytiva). The peak elution fractions were analyzed on a 1% agarose gel stained with SYBR-safe. Fractions containing the plasmid library were pooled and precipitated with ethanol. The final plasmid library was resuspended in nuclease-free H_2_O to a final concentration of 100 ng/µl, aliquoted, and stored at -20^°^C.

### Emulsion PCR

Emulsion PCR was performed based on a previous described procedure ^117^. To amplify the cloned library to produce the linear DNA templates for GATO-seq. A PCR master mix was prepared on ice, consisting of 1X HF Phusion buffer (New England Biolabs), 3.2 µM forward (5′-CAC TCC ACT CGG AGC CAG CA-3′) and reverse (5′-GCG GTG GAG CTC CAA TTC G -3′) primers, 1.6 ng/µL plasmid library, and 2 mM dNTP mix in a total volume of 2 mL. Phusion DNA polymerase was added at a final concentration of 2 units/µL, and the reaction was kept on ice.

The emulsion was generated by mixing 17.8 mL of ice-cold mineral oil (Sigma), 4.6 mL ABIL-90 (Evonik), and 10 µL Triton X-100 (Sigma), followed by vortexing at maximum speed for one minute. After allowing the mixture to settle on ice, the PCR master mix was gradually added dropwise to the vortexing oil phase. The emulsion was stabilized by an additional vortexing step at maximum speed for five minutes at room temperature. The emulsion was then aliquoted into a 96-well PCR plate (100 µL per well) and subjected to thermal cycling for 30 cycles using an annealing step of 10 s at 62^°^C and an extension step for 30 s at 72^°^C.

Following amplification, the PCR products were recovered by centrifugation at 10,000 × g for 10 minutes at 4°C. The lighter oil phase was aspirated, and residual oil was removed through sequential washes with water-saturated ethyl ether and ethyl acetate (Sigma). The final aqueous phase, containing the PCR product, was carefully extracted to avoid contamination from the gel-like interphase. Ethyl acetate was further removed by an additional ether wash, followed by drying in a SpeedVac concentrator for 10 min.

The DNA was precipitated by adding two volumes of 100% ice-cold ethanol and incubated at -20°C overnight or at -80°C for one hour. The precipitate was collected by centrifugation at 10,000 × g for 30 minutes at 4°C, washed with 70% ethanol, air-dried, and resuspended in nuclease-free water. The purified PCR product was applied to a 1 mL HiTrap Q column equilibrated in TE buffer supplemented with 400 mM NaCl. After washing with 10 column volumes, the product was eluted using a linear gradient of 400 to 600 mM NaCl over 20 column volumes. The purity of the eluted fractions was assessed by agarose gel electrophoresis. Fractions containing the PCR product were precipitated with ice-cold ethanol as described above and resuspended to a final concentration of 10 µg/µL in nuclease-free H_2_O, aliquoted, and stored at -20^°^C.

### RNA spike-in preparation

The custom RNA spike-in was designed to cover the expected RNA size products from GATO-seq experiments. 10-, 18-, 50-, and 100-mer sequences were chosen from a random sequence generator (https://faculty.ucr.edu/~mmaduro/random.htm) with GC content adjusted to 0.7 to match the library composition. The sequences were confirmed to show no overlap with the reference GATO-seq library by BLAST and sequence alignment. The 5’ adapter sequence, comprising the 9 nt G-less cassette and 10 nt cloning adapter were added to each RNA to match the level of redundancy of the transcript from GATO-seq experiments. RNA molecules were purchased from Sigma-Aldrich with the following sequences corresponding to the 10-mer: 5’-acu cca cuc gga gcc agc aGA ACC CAC UC-3’, 18-mer: 5’ - acu cca cuc gga gcc agc aUA GUC GUC AUA UAU AGA U-3’, 50-mer: 5’- acu cca cuc gga gcc agc aGG GAC UCA UCU GCA GAG CUA UAG CGU GGG AAA CAA UUC CGC CGC GG AUU C-3’, 100-mer: 5’-act cca cuc gga gcc agc aGG CAG CGC CUC AGA GUA AGU AAU AGA CUA AAG GAA GGC UCA UAU ACU CGA CAG UCG AGG CGC GGU AAC CAA GUU AGU CGC GGU GUC CGU AGC CGC UCU UC-3’. Lower case sequences indicate the common 5’ cloning adapter. RNAs were resuspended in nuclease-free H_2_O, aliquoted and stored at -80 ^°^C.

For the 300-mer, a portion of the human LINE-1 ORF-2 coding region was cloned into a vector containing a T7 promoter and 5’ adaptor sequence and a 5’ hepatitis delta virus (HDV) ribozyme. The fragment comprising the T7 promoter and template DNA was amplified by PCR in a 10 mL reaction. The PCR product was purified via anion exchange, using a 1 mL HiTrap Q column (Cytiva) equilibrated with 20 mM Tris-HCl pH 7.5, 10 mM EDTA pH 8.0 and 400 mM NaCl. The PCR product was eluted with a gradient up to 600 mM NaCl in equilibration buffer over 20 column volumes. The elution peak was concentrated by ethanol precipitation, and the PCR product was used directly for T7 RNA polymerase in vitro transcription reactions. Reactions were carried out in buffer containing 40 mM Tris-HCl pH 7.9, 6 mM MgCl_2_, 10 mM DTT, 10 mM NaCl, 0.001% Triton X-100, and 2 mM spermidine in the presence of 4 mM each ribonucleotide, 0.9 µM T7 RNA polymerase and 100 µg of template DNA. The reaction was incubated for 3h at 37 ^°^C and quenched with 70 mM EDTA and 300 mM NaCl. The nucleic acids were concentrated by isopropanol precipitation, and the RNA transcript was purified by gel extraction from a 6% polyacrylamide TBE-urea gel. The full reaction was loaded on the gel, and the band position was determined by TLC UV-shadowing, excised, and crushed by passing it through a syringe. The RNA was eluted from the polyacrylamide by soaking the crushed gel band in 300 mM sodium acetate pH 5.3 (Sigma Aldrich) and incubated for 10 min at 75^°^C and shaking at 700 rpm. Small acrylamide pieces were removed using a Costar Spin-X centrifuge tube (Corning). Finally, the RNA was purified and concentrated using a Monarch RNA cleanup kit (100 µg) (New England Biolabs). RNA concentration was estimated using the Qubit RNA high sensitivity detection kit (Thermo Scientific), aliquoted, and stored at -80^°^C. RNA integrity was assessed by polyacrylamide TBE-urea gel electrophoresis and RNA fragment analysis. The final 300-mer consisted in the following sequence: 5’-acu cca cuc gga gcc agc aGU UCA GGA AAC CCA UCU CAC GUG CAG AGA CAC ACA UAG GCU CAA AAU AAA AGG AUG GAG GAA GAU CUA CCA AGC CAA UGG AAA ACA AAA AAA GGC AGG GGU UGC AAU CCU AGU CUC UGA UAA AAC AGA CUU UAA ACC AAC AAA GAU CAA AAG AGA CAA AGA AGG CCA UUA CAU AAU GGU AAA GGG AUC AAU UCA ACA AGA GGA GCU AAC UAU CCU AAA UAU UAU GCA CC CAA UAC AGG AGC ACC CAG AUU CAU AAA GCA AGU CCT CAG UGA CCU ACA AAG AGA CUU AGA CUC CCA CAC A-3’

### GATO-Seq Sample Preparation

Preinitiation complexes (PICs) were assembled in a stepwise manner as described ^114^. Reactions (250 µL final volume) at the following final buffer composition: 20 mM Tris-HCl pH 8.0, 100 mM NaCl, 4% (v/v) glycerol, 2 mM MgCl_2_, 1 mM DTT, 25 µM ZnCl_2_. *S. scrofa* RNA polymerase II was assayed at 100 nM final concentration and DNA template at 112.5 nM. Initiation factors were included at the following molar ratios relative to RNA polymerase II: TFIIH/CAK (1:1), TFIIA (1:5), TFIIB (1:2.5), TBP (1:2.5), TFIIE (1:1), and TFIIF (1:5).

The TFIIH core was preincubated with CAK at 30°C with shaking at 300 rpm. The upstream complex was formed by incubating template DNA with TFIIA, TFIIB, and TBP under the same conditions. RNA polymerase II was preincubated with TFIIF at 30°C and 300 rpm. TFIIE was added to the TFIIH/CAK complex and incubated for 5 min. The upstream complex was then combined with the RNA polymerase II/TFIIF complex, followed by the addition of the TFIIH/CAK/TFIIE mixture. The reaction was brought to the final volume with a 4X reaction buffer and nuclease-free water and incubated at 30°C for 20 min at 300 rpm. Initial transcribing complex and G-less cassette were done simultaneously by adding ATP to a final concentration of 1 mM (necessary for TFIIH helicase and CAK kinase activities), and CTP and UTP to a final concentration of 10 µM, or 1 mM where indicated, followed by a 10 min incubation at 30^°^C, 300 rpm. In reactions where elongation factors DSIF and NELF were included, they were added in 1:3 molar ratio relative to final RNA polymerase II concentration and incubated for additional 10 min at 30^°^C and 300 rpm. Following the incubation, triptolide (Sigma-Aldrich) was added to a final concentration of 10 µM. GTP was immediately added to a final concentration of 10 µM (or 1 mM where indicated) to start the reaction, and samples were incubated at 30°C and 300 rpm. For experiments using 1 mM NTP, 1 mM ATP was added again together with GTP to compensate depletion during initiation reactions. For experiments including TFIIS or TFIIS DE 282-283AA mutant, these factors were added at this step.

Transcription reactions were quenched by adding an equal volume of phenol:chloroform:isoamyl alcohol (25:24:1) (Thermo Fisher), 0.1 pmol RNA spike-in was added for calibration. The sample was vortexed for 10-30 s and centrifuged at 15,000 rpm for 5 min. The aqueous phase was transferred to a new tube. One volume of chloroform was added, vortexed, and centrifuged again under the same conditions. The upper phase was recovered, mixed with 1/10 volume of 3 M sodium acetate pH 5.2 and three volumes of 100% ethanol, and incubated at -20°C for at least 1 h. RNA was pelleted by centrifugation at 15,000 rpm for 30 min at 4°C, washed with 70% ethanol, and air-dried for 2 min before resuspension in 170 µL nuclease-free water. The residual DNA template was removed (sample volume 170 µL) by mixing the sample with 20 µL of 10X Turbo DNase buffer and 10 µL of Turbo DNase (20 units) at 37°C for 1 h (Thermo Fisher). The reaction was purified using the Monarch RNA Cleanup Kit (100µg) (New England Biolabs) following the manufacturer’s protocol and eluted in 20 µL nuclease-free H_2_O. RNA products were polyadenylated using *E. coli* poly(A) polymerase (New England Biolabs). RNA was combined with 4 µL 10 X Poly(A) polymerase reaction buffer, 4 µL of 10 mM ATP (1 mM final) and 10 µL nuclease-free H_2_O. The reaction was initiated by adding 2 µL poly(A) polymerase and incubated at 37°C for 5 min in a thermocycler. The reaction was quenched with 10 µL 50 mM EDTA pH 8.0 (10 mM final) and placed on ice. RNA was purified using the NEB Monarch RNA Cleanup Kit and eluted in 11 µL nuclease-free H_2_O. Final RNA concentration was measured using a Qubit RNA High Sensitivity Assay Kit (Thermo Scientific). Yields ranged from 30-100 ng. Samples were either flash-frozen in liquid nitrogen and stored at - 80°C or used immediately for Oxford nanopore direct RNA sequencing library preparation.

### GATO-seq nanopore library preparation and sequencing

Sequencing libraries were prepared with SQK-RNA004 Direct RNA sequencing kit (Oxford Nanopore Technologies) following the protocol provided by the manufacturer, with the following modifications: Total pure NGS cleanup magnetic beads (Omega Biotek) were used instead of AMPure XP beads for all adaptor excess removal steps and Superscript IV reverse transcriptase (Thermo Fisher) or Induro (New England Biolabs) were used for first strand synthesis. Direct RNA libraries were loaded in FLOMIN004RA sequencing flow cells (Oxford Nanopore Technologies) and sequenced on a MinION sequencing device (Oxford Nanopore Technologies) operated with the latest available version of MinKNOW software (v23.07 and above) (Oxford Nanopore Technologies) for MacOS. Sequencing reactions were stopped after ∼1,000,000 raw reads were acquired (3-7 h). Flow cells were washed with EXP-WSH004-XL (Oxford Nanopore Technologies) following the manufacturer’s instructions and reused if they displayed at least ∼800 pores available for sequencing using the Flow Cell checking options in MinKNOW.

### Original code

Original code reported for this manuscript was written for Python and initially generated by ChatGPT (V4 to 5) except for the sequence retrieving code, written in Jupyter Notebook (v7.4.7). Generated code was examined and manually tested to corroborate functionality before being incorporated to the analysis. All custom Python scripts are available in Zenodo, unless otherwise indicated.

### GATO-seq data analysis

POD5 files generated during direct RNA sequencing were basecalled using Dorado 9 (Oxford Nanopore Technologies) with the super high accuracy model (rna004_130bps_sup@v5.1.0) and with a minimal threshold Phred score set to 7. The resulting FASTQ files were aligned to the reference spike-in sequences for calibration and to the GATO-seq promoter proximal sequences using Bowtie2 (v2.4.5)^118^, with poly-A tails soft-clipped during alignment. To generate the reference sequence file, sequences of the 1000 gene fragments present in the template library were catenated. The sequences were separated by 100bp poly-T spacer sequences to reduce overlapping noise during alignment. The reference FASTA, bed file with coordinates of gene-specific TSS and end of transcript, as well as GFF annotation files are available at (GEO GSE309328).

The shortest RNA spike-in (10 nt) was unmappable, and the shortest unambiguously mapped RNA corresponded to the 18-mer. The number of reads associated with spike-in RNAs displayed a proportional relationship with RNA length, which was used to normalize length-dependent sequencing bias: the absolute number of 3’ends within a window of 5 nt of each RNA from the spike-in was obtained using deepTools (v3.5.5)^119^ bamCoverage with the --offset -5 option. These counts were normalized relative to the total number of detected 3’ ends. The normalized 3’ end counts were plotted as a function of RNA length and fitted to a straight line. A theoretical even distribution of reads across the four equimolar RNA species corresponds to a value of 0.25. To correct for length bias, we calculated a length-specific correction factor to adjust the observed slope from the linear regression of the spike-in counts to the theoretical 0 slope with origin at 0.25. The estimation of the correction factors was done in Excel (Microsoft).

For gene library-aligned reads, counts were first normalized using counts per million (CPM) to account for sequencing depth differences and leveraging the fact that all gene templates are equally sized. Only 3’ ends were retained using the --offset -1 option in deepTools (v3.5.5). The length correction factor derived from the spike-in analysis was then applied to the CPM-normalized bedgraph files using a custom Python script. Replicate consistency was assessed by PCA analysis and correlation between replicates in deepTools (v3.5.5).

Consistent replicates were averaged to generate signal profiles in deepTools (v3.5.5) using bigwigAverage function. Per-base passthrough indices were estimated from these normalized bedgraph files using a custom Python script. The script estimates the upstream signal + α/total signal + 2α), where α=1 and is included to avoid impossible mathematical outcomes. The script generates a bedgraph with passthrough index-converted at each position. Genes without signal were excluded from this calculation. These bedgraph files were then used to generate the cumulative distribution graphs using the computeMatrix function in deepTools (v3.5.5). The cumulative distribution midpoint was estimated from the position at y=0.5. Cumulative distribution midpoint vs time plots were fit to a straight line using a linear regression in GraphPad Prism (V10).

To identify and compare pause sites, the signal of each gene was normalized to the total signal within each gene using a custom Python script, sites with-out signal were assigned a zero value. Then, the average and the standard deviation of the signal within each gene from the bedgraph files were calculated. Pause site coordinates were extracted for positions where signal was at least four standard deviations above the average signal and passed this threshold in at least two consecutive time-points. Sites from the last 50 bp were omitted from the final list of pause sites to avoid accounting for full length transcripts. These calculations were achieved using a custom Python script, which generated an output bed file with pause sites coordinates, signal and z-score values from each time point (z=GATO-seq signal - gene average/gene standard deviation) To identify pause sites that display exponential, time-dependent decay, a Python script was used. The script fits the semi log fraction of GATO-seq signal vs time plots to a straight line and identifies sites where the 5-min time point showed the lowest signal and intersects with the 1 minute time point. A dynamic residual range set at a 0.1 threshold (10%) was used to estimate the goodness of fit. y-intercept and slope were used to calculate the E_max_ and t_1/2_, respectively.

To detect changes upon TFIIS treatment, pause sites were defined using the procedure described above. For this set of experiments, only 1- and 5-minute time points were available. Therefore, pause sites must pass the 4-standard deviation threshold in both time points. Linear fit was not attempted as only two time-points were available. Pause positions between the two datasets were compared to identify overlaps with a tolerance of ±3 nt. Bed files containing the coordinates of overlapping and non-overlapping positions between datasets were generated. Overlapping sites were further classified based on the signal change to the TFIIS condition using --kmeans clustering restricted to the TFIIS 1 minute dataset using deepTools (v3.5.5). The number of clusters was determined empirically until clusters became redundant (3 clusters).

The output bed files containing coordinates of TFIIS-sensitive -emergent and the 3 clusters of the shared pauses were used to retrieve the surrounding nucleotide composition using a custom Python script. Sequence logos were generated using kpLogo: k-mer probability logo^120^, with the GATO-seq library provided for background estimation. (http://kplogo.wi.mit.edu/index.html?).

### Sequence scanning search and analysis

A text file containing a list with 43 sequences classified as super pauses (super pause sequences), each sequence consisting of 16 bp (12 bp upstream and 3 bp downstream flanking the pause position), was generated and used to scan 10,000 promoter proximal regions from protein coding genes for hypothetical super pause sites. Coordinates of TSS were obtained from the GetGenAnnotation pipeline (https://zenodo.org/records/5519928) ^116^ from HEK293T (same dataset used for GATO-seq library gene selection and annotation) and used to generate a bed file with coordinates from human genome hg38.14 assembly from +1 to +300 bp. The bed file and the hg38.14 FASTA human genome assemble were used to generate a FASTA file containing only the promoter proximal sequences using the getfasta function in Bedtools (v2.31.1)^121^. Scanning was performed using a custom Python script, which scans each sequence with a 1 bp step size. Background control sequences were generated using *fasta-shuffle-letters* (v5.4.1) from the MEME suite^122^, using the -k 2 option to preserve the di-nucleotide composition.

The resulting bed file containing coordinates, sequences and relative positions of hypothetical super pause sites, and background sequences was used for further analysis. Alternatively, motif search was done using FIMO (v 5.4.1) from MEME suite^123^, providing a position weight matrix as a query, generated from the relative nucleotide frequencies from the 43 super pause sequences, and using a first-order Markov model to account for sequence background composition. FIMO searches were unable to detect our known super pause sites, and was thus not pursued further to find additional hypothetical super pause sequences.

Gene ontology analysis was done in GOrilla^124^ (https://cbl-gorilla.cs.technion.ac.il/) providing the 10,000 protein coding genes as a background list and the list of genes with hypothetical super pause sites as a query, and with all other settings as default.

Gene coordinates for the hypothetical super pause sites were used to obtain the PRO-seq and ChIP-exo signals. PRO-seq signal was obtained from^92^ GSM4730174 (datasets GSM4730174 and GSM4730175) and ChIP-exo signal was obtained from ^91^ GSM6703891(sample GSM6703891). Heatmaps and profile graphs, were generated using deepTools (v3.5.5).

### Cryo-EM sample preparation

The RNA polymerase II complex was assembled as described above for the RNA extension assay on the Type III “Super pause” consensus scaffold. 100 pmol final RNA polymerase II was mixed with 125 pmol RNA-DNA template, followed by 125 pmol non-template DNA. In contrast to RNA extension assays, the transcription complex was at a 10-fold higher concentration for cryo-EM preparations to ensure adequate number of particles on grids. The final buffer condition consisted of 100 mM NaCl, 20 mM Na-HEPES pH 7.4, 3 mM MgCl_2_, 1 mM DTT, and 4% (v/v) glycerol. The sample was incubated for 30 min at 30°C. RNA extension was enabled by adding CTP, GTP and UTP to a final concentration of 10 µM and incubated for 5 min at 30 °C. Extension was quenched with 10 mM EDTA pH 8.0 (final concentration) and transferred to a 4 °C ice bath. The complex was purified by applying it to a Superose 6 increase 3.2/300 column equilibrated in buffer matching reaction conditions (100 mM NaCl, 20 mM Na-HEPES pH 7.4, 4% (v/v) glycerol, 3 mM MgCl2, and 1 mM DTT) on an Äkta Micro purification system (Cytiva) at 4°C. 50 µL fractions were collected. Peak fractions were analyzed by SDS-PAGE followed by Coomassie staining and TBE-urea gel electrophoresis.

The peak fractions containing RNA polymerase II and nucleic acids were crosslinked with a final concentration of 0.1% (v/v) glutaraldehyde for 10 min on ice and quenched with 8 mM aspartate and 2 mM lysine. The crosslinked sample was dialyzed for 4 hours at 4°C against a buffer containing 100 mM NaCl, 20 mM Na-HEPES pH 7.4, 1 mM DTT, and 3 mM MgCl_2_, in 10 kDa MWCO Slide-A-Lyzer MINI Dialysis Units (Thermo-Fisher). Dialyzed samples at a final concentration of 300-400 nM were applied to R2/2 UltrAuFoil grids (Quantifoil). The grids were glow-discharged for 2 min before applying 2 µL of sample to each side of the grid (4 µL total), incubated for 4 s, and blotted for 4s. Vitrification was done by plunging the grids into liquid ethane using a Vitrobot Mark IV (FEI Company) operated at 4°C and 100% humidity.

### Cryo-EM data collection and processing

Cryo-EM micrographs were collected on a FEI Titan Krios II transmission electron microscope operated at 300 keV. A K3 summit direct detector (Gatan) with a GIF quantum energy filter (Gatan) was operated with a slit width of 20 eV. Data was automatically acquired using FEI EPU software at a nominal magnification of 105,000×, corresponding to a pixel size of 0.822 Å per pixel. Image stacks of 50 frames were collected over 1.53s in counting mode. The dose rate was 22.54 e^−^/pix/s for a total dose of 50.5 e^−^ per Å^2^. A total of 9,160 image stacks were collected.

Frames were stacked, contrast-transfer-function was estimated, and motion-correction was applied in CryoSPARC live v4.7.0 ^125^. Particles were auto picked using the blob picker tool in CryoSPARC live v4.7.0 and extracted using a box size of 400 pixels. Particles were used for 2D classification and classes where secondary structure features were observed were used for ab-initio 3D reconstruction in CryoSPARC v4.7.0. Non-aligning particles were classified and discarded by 3D classification with image alignment (Heterogeneous refinement), using the ab-initio volume as an input reference.

An initial volume of RNA polymerase II at 2.8 Å nominal resolution was generated using homogenous refinement. Particles that displayed a flexible clamp domain were removed using 3D classification without alignment at a 5Å initial resolution. A solvent mask that encompassed RNA polymerase II was generated from a previously published RNA polymerase II-TFIIS model (PDB 8A40). TFIIS atoms were removed before generating a 15Å low-pass filtered volume in ChimeraX (v1.7) ^126^, used as an input for mask creation in CryoSPARC v4.7.0 with a dilation radius and soft-padding of 12 and 5 pixels respectively. Particles with clear clamp density were classified further at an initial 4Å resolution using an active site mask comprising RPB1 bridge helix residues, 844-860, trigger loop residues 1083-1103 and 3 base pairs of the RNA-DNA hybrid. A 10Å low-pass filtered volume was created in ChimeraX (v1.7) and a mask was produced in CryoSPARC v4.7.0 with 3 pixels dilation radius and 1 pixel soft padding. Ab-initio reconstructions were done from particles of each class, followed by homogenous refinement. The sidetracked conformation emerged from this classification, whereas pre- and posttranslocated states required further rounds of classification. Other 3D classes showed weak DNA-RNA hybrids and further local refinement did not yield improvement. The maps showed some orientation bias. To alleviate this, the rebalance orientations job was implemented in CryoSPARC v4.7.0. The final sidetracked map was improved by using reference-based motion correction and providing the refined sidetracked volume as an input, followed by global CTF correction and homogenous refinement. Pre- and posttranslocated maps were obtained after 3D classification at an initial 3Å resolution, masking 3 base pairs upstream of the 3’ end of the RNA-DNA hybrid on the orientation rebalanced and reference-based motion corrected input volume. A mask was generated by providing a 3Å low-pass filtered volume into CryoSPARC v4.7.0 with 1-pixel of dilation radius and soft padding. Classified particles were used for ab-initio reconstructions, followed by global CTF correction and homogenous refinement. Maps were sharpened using a B-factor of -20 Å^2^. Resolutions were estimated by 0.5 FSC threshold in CryoSPARC v4.7.0.

### Model building

RNA polymerase chains 1-12, and nucleic acids from PDB 8A40 were rigid body fit in the refined cryo-EM maps in UCSF ChimeraX (v1.7). Side chains were manually adjusted in Coot 1.1.08 ^127^. Densities for RPB4 and RPB7 (stalk) were weak and were not adjusted after the rigid body fit. Trigger loop residues 1104-1116 and 1105-1116 were not visible in the sidetracked and pretranslocated maps, respectively and therefore were not included in the final models. The resolution of the active site permitted unambiguous assignment of the sequence register. The nucleic acids from PDB 8A40 ^81^ were substituted for the corresponding bases in the Super Pause consensus sequence, and the RNA 3’ end (1 nt for pretranslocated and 2 nt for sidetracked) were added to the corresponding models and manually adjusted. We modeled 13 bp of the downstream DNA duplex and 10 bp of the RNA-DNA hybrid. Density for the 12 deoxynucleotides of the non-template DNA strand were not visible and therefore not included in the final models. Upstream DNA densities were weak, and commonly this region requires focus refinement and compose-map creation, we therefore opted for only rigid body fitting 8 bp of the DNA duplex.

Final models were obtained by real space refinement in Phenix version 1.21.2 ^128^ with secondary structure and Ramachandran restrains. PDB input models for refinement were prepared using ReadySet. Statistics about data collection and modelling are in Data Table 1.

