## Supplementary figures for "A sequence-encoded promoter proximal super pause stabilizes an offline RNA polymerase II state"

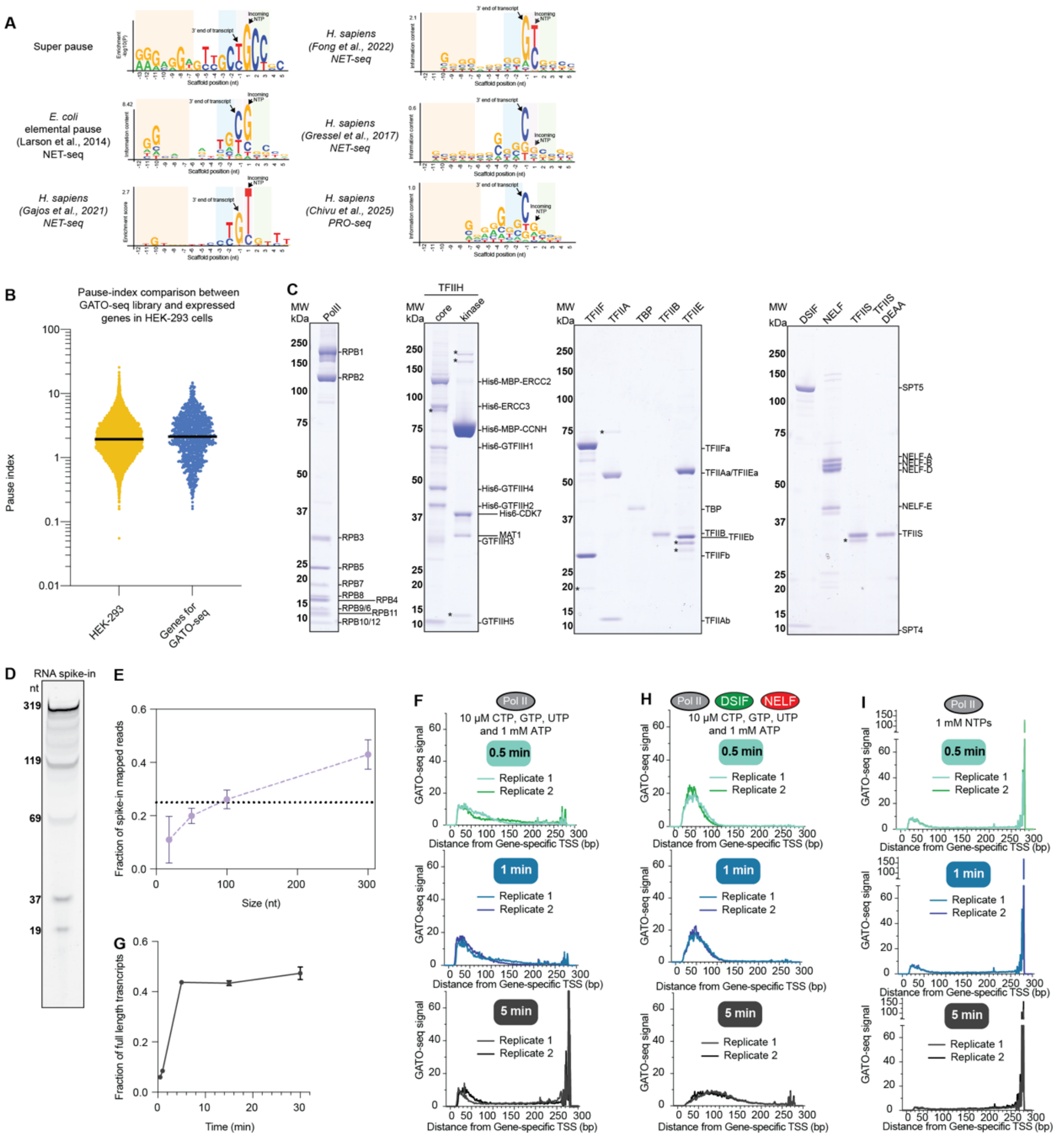

Supplementary Fig. 1

**Supplementary Figure 1. Library comparison, protein preparation, and quality control of GATO-seq.**

**(A)** Comparison of super pause consensus sequence to previously reported pause sequence logos[45,46,48,52,97]. (**B**) Pause index comparison between 10,000 protein coding genes in HEK293T cells and the 1,000 gene subset of these genes used for GATO-seq library preparation. Pause index = PRO-seq 3'end counts from TSS to + 100 bp / PRO-seq 3'end counts from 150 to + 300 bp. Yellow dots correspond to all 10,000 protein coding genes (**Supplementary Table 1**) and blue dots correspond to the subset selected for GATO-seq templates. (**C**) Coomassie stained SDS-PAGE (4-12%) gels of purified proteins. (**D**) RNA spike-ins used for calibration of GATO-seq experiments visualized by SYBR-gold staining on 10% polyacrylamide, 6 M urea gel. (**E**) Normalized fraction of spike-in RNA Oxford-nanopore reads. Mean and standard deviation from three independent replicates. Horizontal dashed line at 0.25 is a reference for the theoretical fraction of a perfectly unbiased sample consisting of equimolar amounts of each spike-in RNA. (**F**) Biological replicates of GATO-seq experiments. Metagene plots generated from the average signal at three different time points of RNA polymerase II with 10 μM NTPs and (**G**) Fraction of full-length transcripts from a 30 minute time course of RNA polymerase II with 10 μM NTPs. Average and standard deviation from two technical replicates. **H-I** Biological replicates of GATO-seq experiments as in (F) with (**H**)DSIF and NELF (**I**) with 1 mM NTPs.

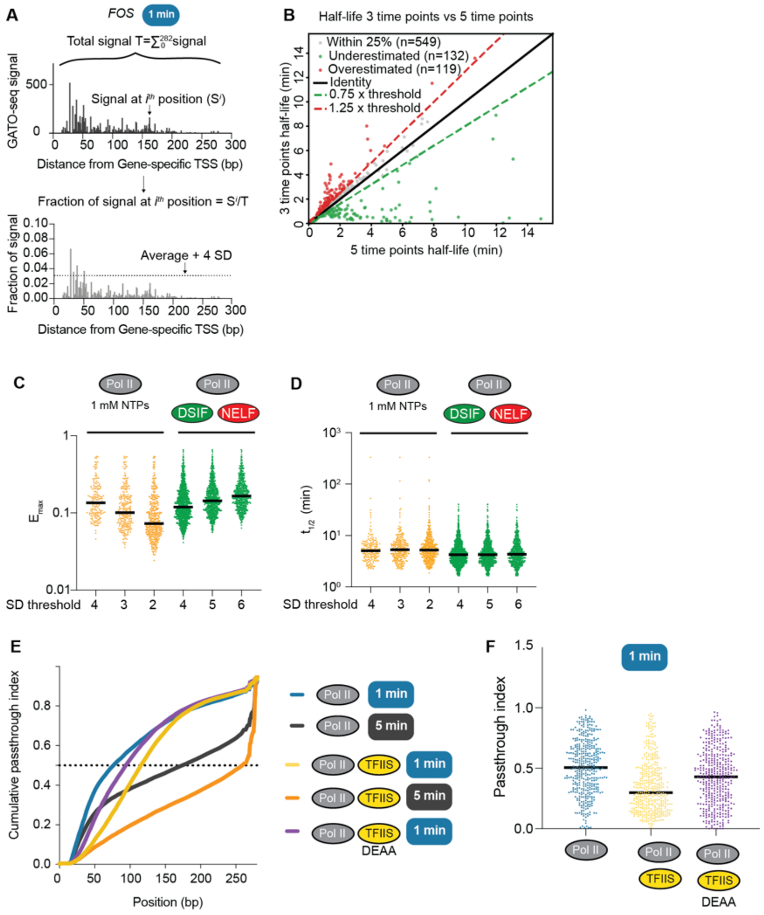


### Supplementary Fig. 2

**Supplementary Figure 2. TFIIS effect on pause behavior.**

**(A)** Schematic of GATO-seq signal normalization to fraction of signal and pause detection threshold**. (B)** Comparison of $t_{1/2}$ values from time courses using 0.5, 1, 5, 15, and 30 minute time points (5 time points, x-axis) against a time course only using 0.5, 1, and 5 min (3 time points, y-axis). Green and red dotted lines indicate 25 % threshold boundaries for underestimated vs overestimated $t_{1/2}$ values. **(C)** $E_{max}$ values estimated from the exponential decay fit of pause sites detected at different standard deviation thresholds for the indicated experimental conditions. (**D**) Half-lives from pause sites detected at different standard deviation thresholds, as in (B) (**E**)

Passthrough index cumulative distributions of indicated conditions. Cumulative density midpoints for TFIIS 1 minute: 118 bp, TFIIS DE282-283AA 1 minute: 97 bp, and TFIIS 5 minutes: 264 bp. (**F**) Passthrough indices at TFIIS sensitive sites from three indicated experimental conditions.

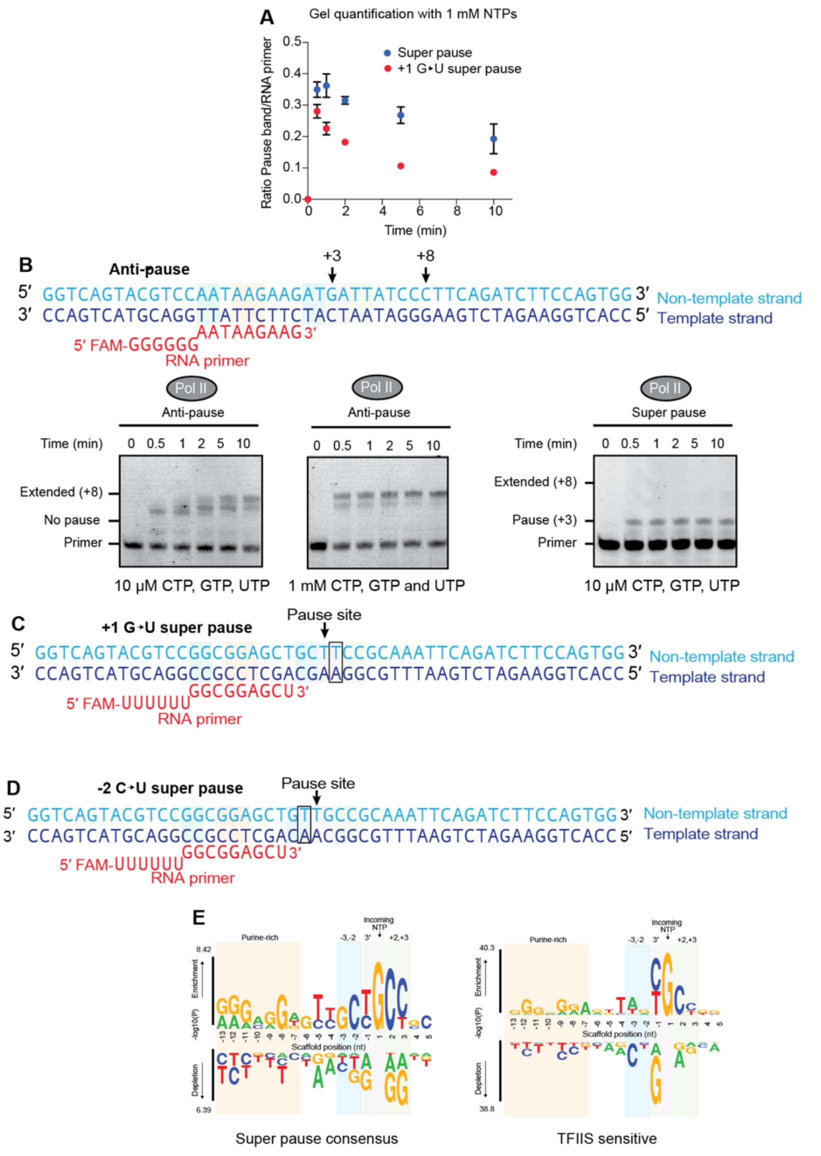


Supplementary Fig. 3

**Supplementary Figure 3. Substitutions to super pause consensus sequence.**

(**A**) Intensity ratio of +3 pause band/ RNA primer quantified from **Figure 4D**. Average and standard deviation from 3 biological replicates. (**B**) Nucleic acid sequence scaffold harboring anti-pause sequence (top) and RNA extension assay on this scaffold (bottom) using 10 μM or 1 mM of indicated NTPs. Super pause RNA extension assay from Fig. 4C is shown on the right for reference. (**C, D**) Nucleic acid sequence scaffolds employed in RNA extension assays, shown in Figure 4 D, E, and F. (**E**) Comparison of super pause logo from Fig. 4A to TFIIS sensitive consensus sequence logo.

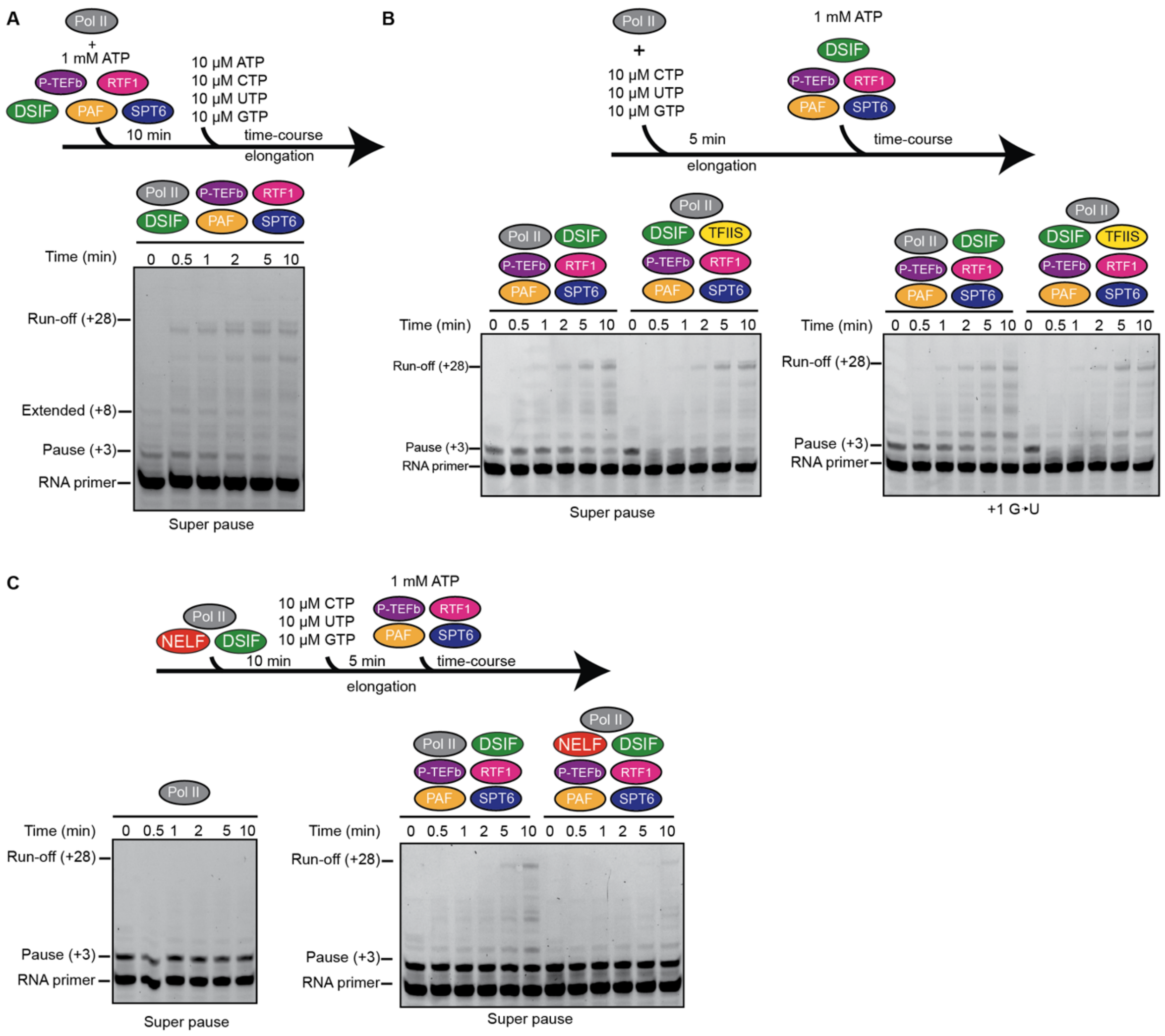


Supplementary Fig. 4

**Supplementary Figure 4. EC* formation on super pause consensus and mutated scaffolds.** (**A**) RNA primer extension time course on super pause consensus sequence scaffold after 10-minute incubation with EC* (1 mM ATP, and 1:3 molar excess of DSIF, P-TEFb, PAF1c (with RTF1) and SPT6 relative to RNA polymerase II) followed by an elongation time course. Due to the addition of ATP during EC* formation, nucleotide misincorporation is observed at time point 0. (**B**) RNA primer extension time course on super pause consensus sequence and +1G to U substituted scaffold as shown in Fig. 5B, in the absence and presence of 1:0.2 molar ratio of

TFIIS. (**C**) RNA primer extension experiments as shown in (B) but pre-incubated with 3-fold molar excess of DSIF and NELF.

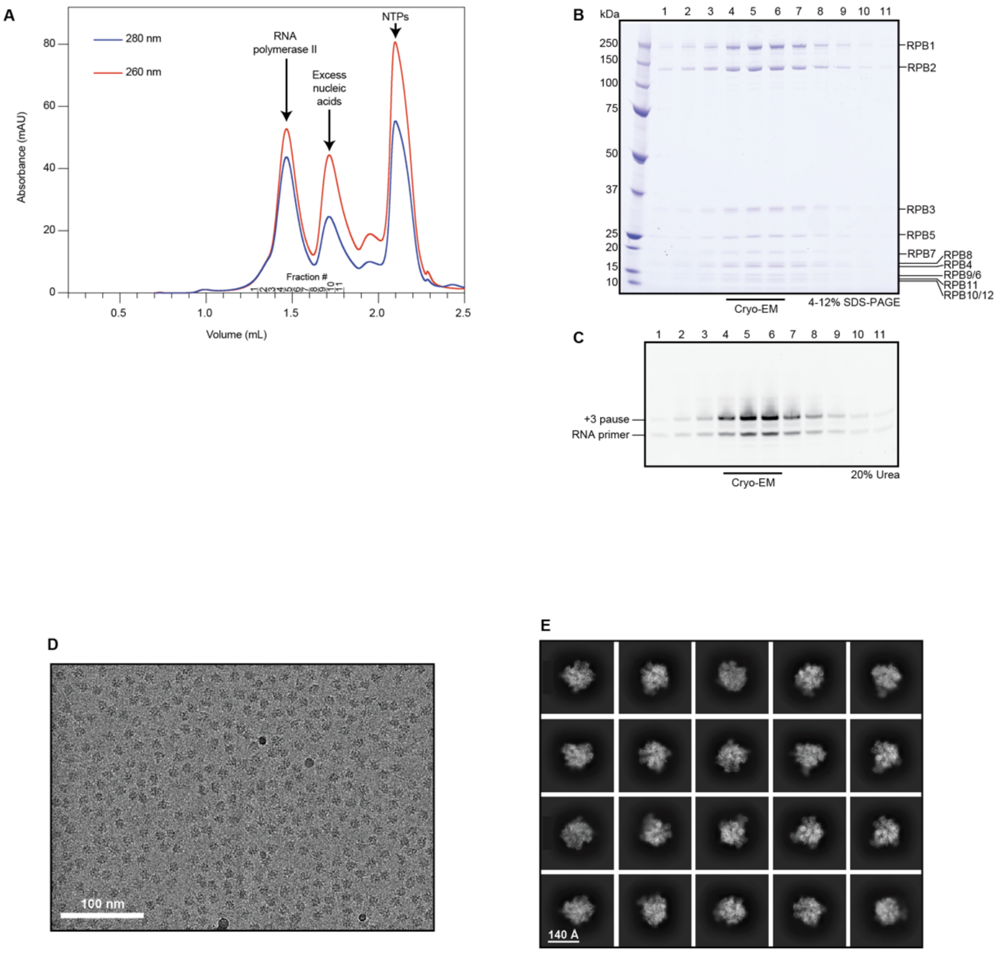


Supplementary Fig. 5

**Supplementary Figure 5. Biochemical preparation of samples for cryo-EM and initial cryo-EM processing.**

(**A**) Size exclusion chromatography traces of transcribing RNA polymerase II on "Super pause" scaffold. (**B**) Coomassie SDS-PAGE of fractions from (a). (**C**) Visualization of RNA extension on a 20% polyacrylamide, 6 M urea gel of fractions from (a) (**D**) Representative micrograph at 2.5 μm defocus. Scale bar shown on bottom left. (**E**) Representative 2D classification images of RNA polymerase II particles. Scale bar is shown on the bottom left.

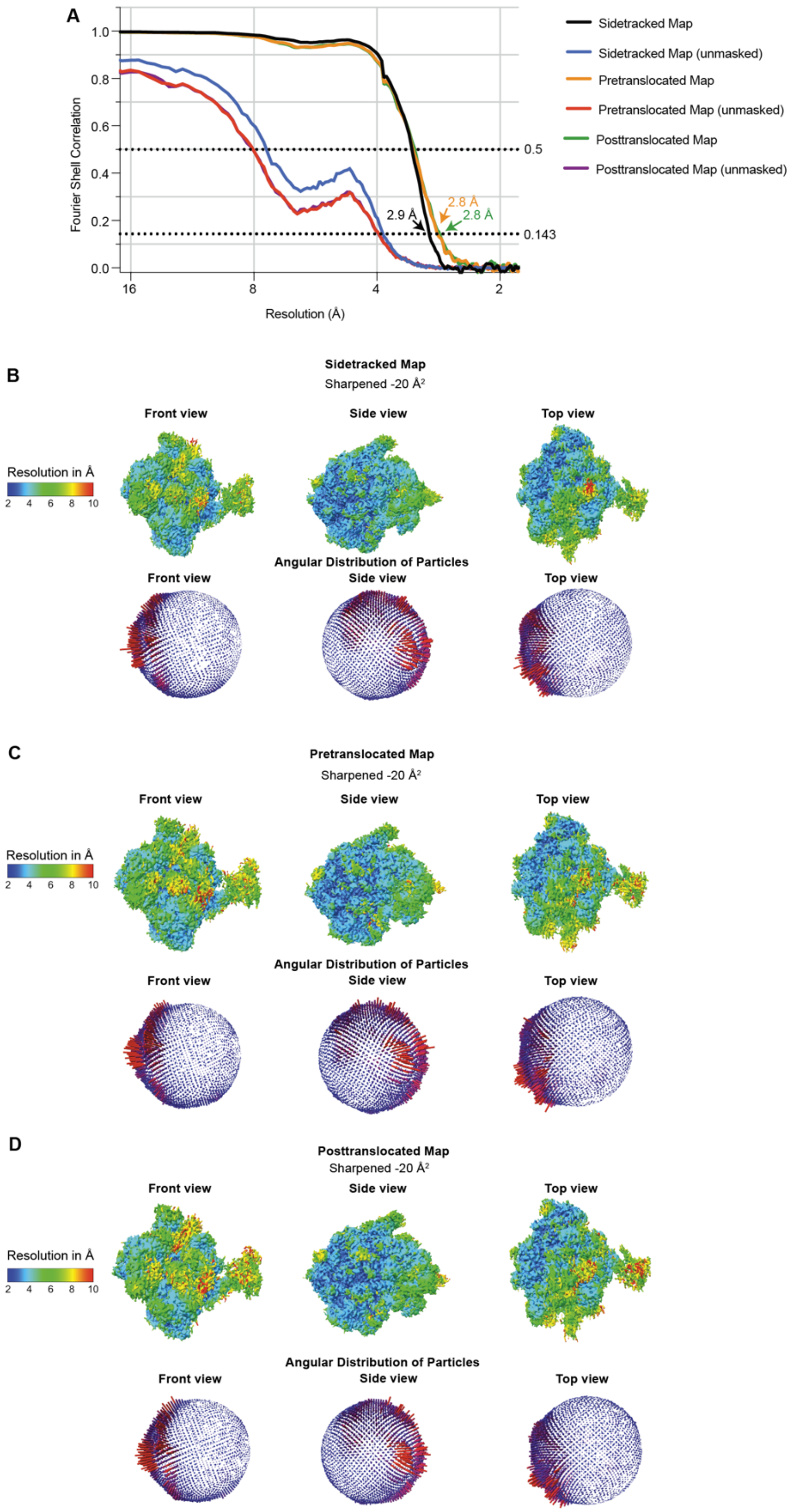


Supplementary Fig. 6

**Supplementary Figure 6. Cryo-EM data quality.**

(**A**) Fourier Shell Correlation curves (corrected and unmasked) of sidetracked, pre and posttranslocated maps. FSC 0.5 and gold standard 0.143 are shown as horizontal dashed lines. Resolution reported for each map is indicated. (**B, C** and **D**) Reconstructions colored by local resolution (Top panels), shading scale shown on left, and angular distribution of particles (bottom panels) from overall refinements for (**B**) sidetracked, (**C**) pretranslocated and (**D**) posttranslocated map.

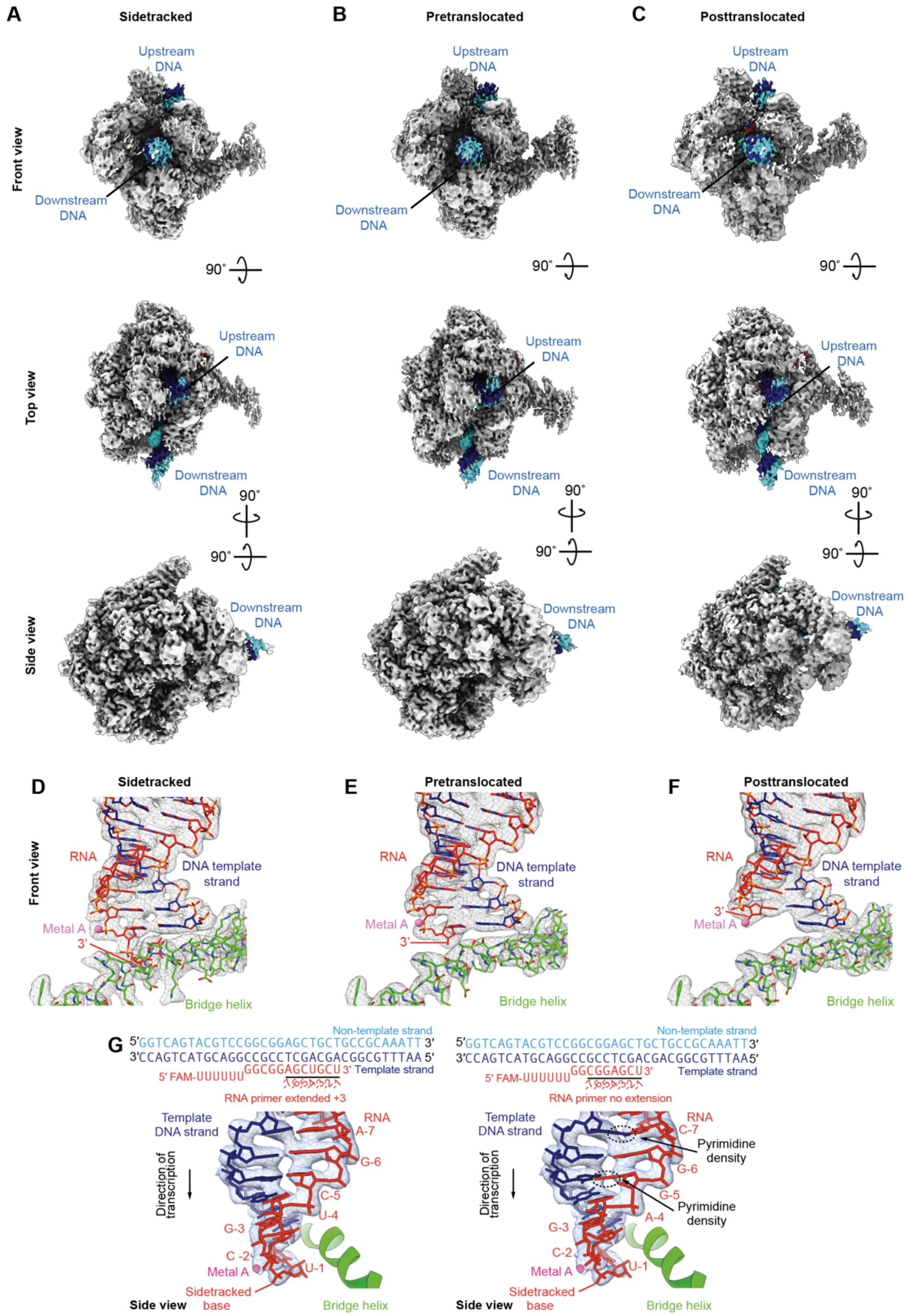


### Supplementary Fig. 7

**Supplementary Figure 7. Overall cryo-EM densities and comparisons of active site translocation states.**

Cryo-EM densities of RNA polymerase in (**A**) sidetracked, (**B**) pretranslocated and (**C**) posttranslocated states. (**D, E** and **F**) Cryo-EM density (in gray mesh) of DNA-RNA hybrid in (**D**) sidetracked, (**E**) pretranslocated and (**F**) posttranslocated (not deposited) states. (**G**) Comparison of DNA-RNA hybrid atomic model fit into cryo-EM density at the super pause (extended +3) on left panel, and without extension on right panel. Guanosine from RNA at position -5 and in the template DNA at position -7 do not fit into the density if RNA extension would have not occurred, whereas all bases at the super pause fit into the density.

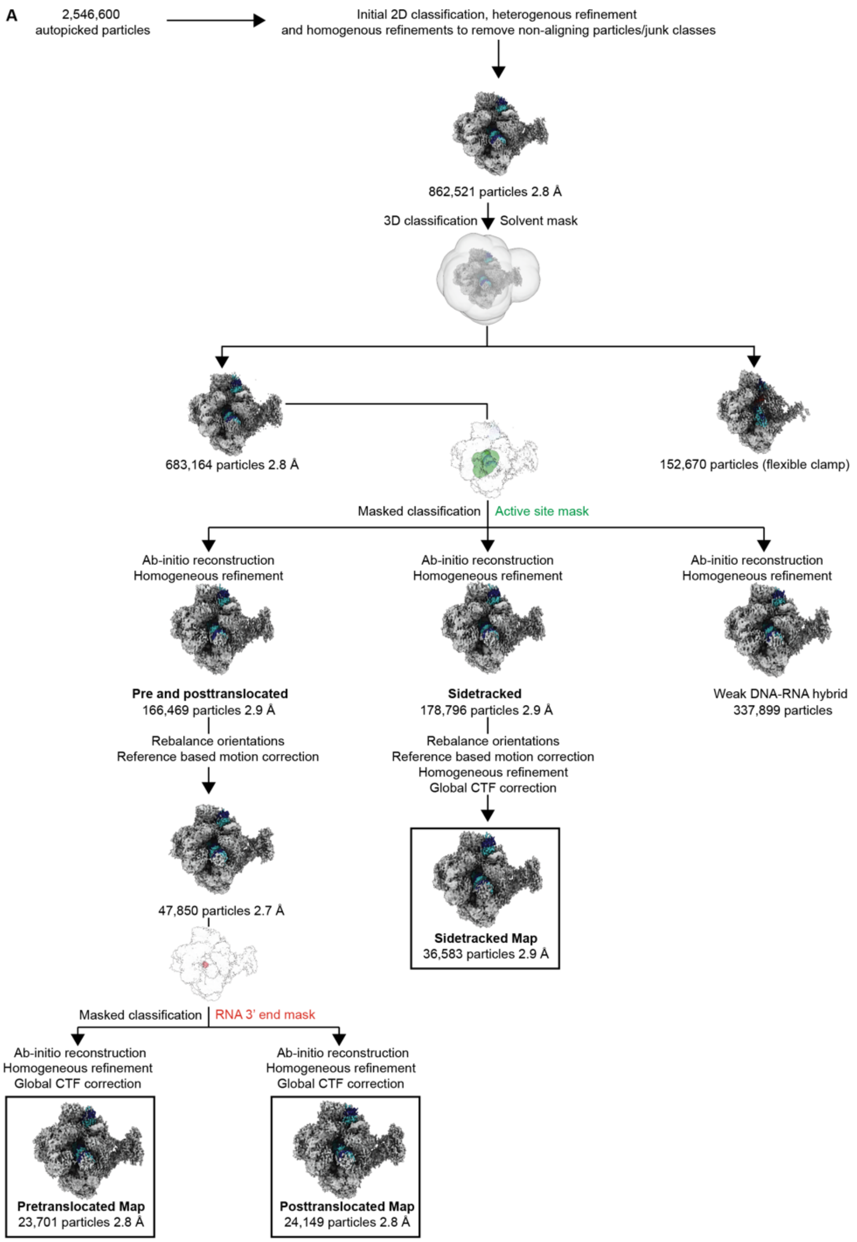


Supplementary Fig. 8

**Supplementary Figure 8. Cryo-EM processing tree for RNA polymerase II on "Super pause" scaffold.**

(**A**) Classification tree for RNA polymerase II on "Super pause" scaffold structures. Resolutions are provided for maps used for further classification. Masks used for focused classification are shown.

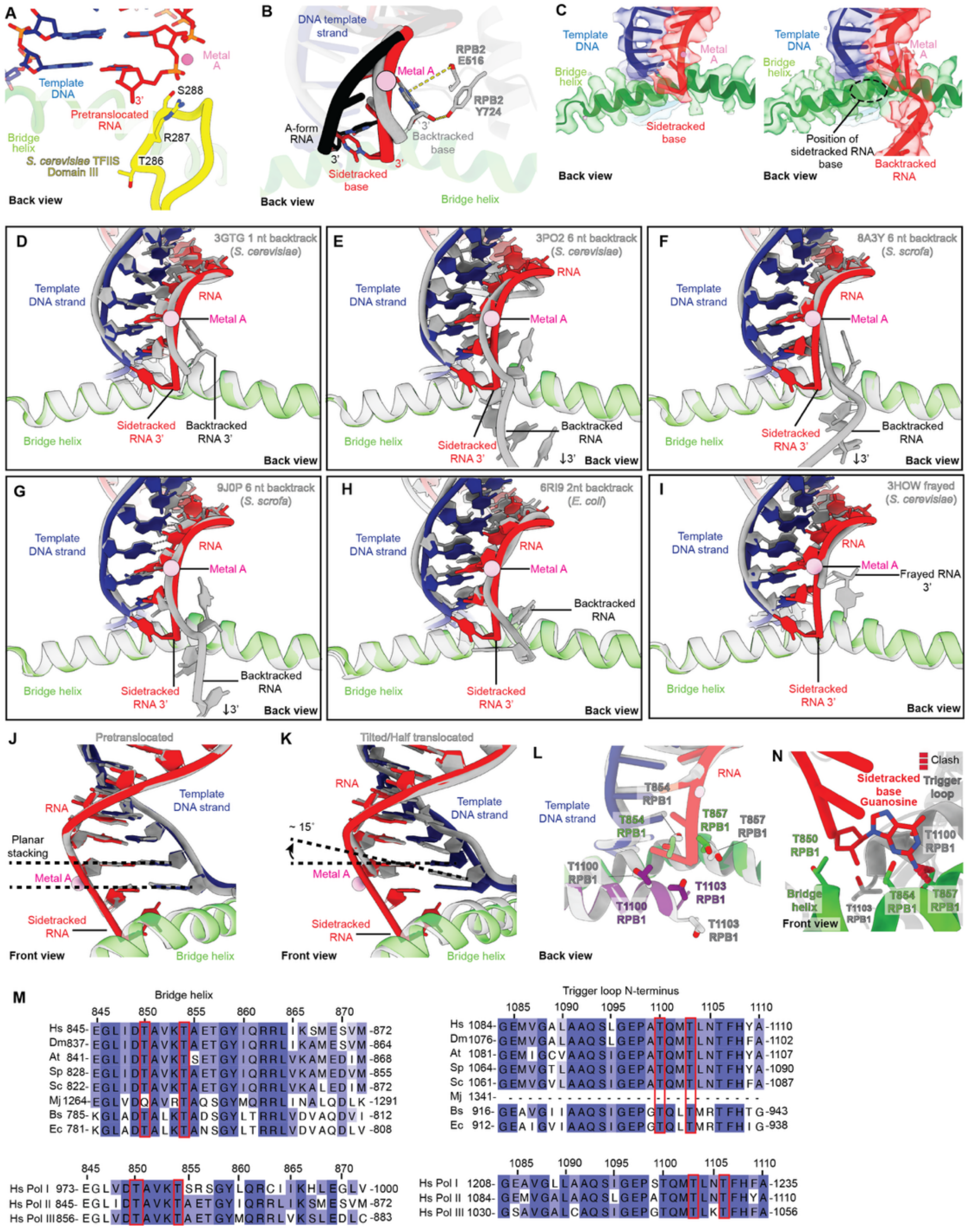

Supplementary Fig. 9

**Supplementary Figure 9. Comparison of sidetracked RNA 3' end to other backtracked and paused conformations**.

(**A)** Overlay of pretranslocated RNA (red) and *S. cerevisiae* TFIIS Domain III (PDB 3PO3) in yellow as in Fig. 6C. No clashes were detected in the overlay. (**B**) Comparison of A-form RNA (black) *S. cerevisiae* 1 nt backtracked RNA (PDB: 3GTG, gray) and *S. scrofa* sidetracked RNA (red). (**C**) Comparison of Cryo-EM density of sidetracked RNA (left) to backtracked RNA from EMD-15127 (right) (**D-I**) RNA polymerase II core alignment between sidetracked RNA (red) and various arrested and paused conformations (gray). PDB entries and species are shown in the upper right corner of each panel. (**J** and **K**) Comparison of register between (J) sidetracked and pretranslocated and (K) sidetracked and tilted/half-translocated RNA-DNA hybrids. (**L**) Back view overlay of sidetracked and activated elongation complex (PDB 6TED) in gray. Threonine pocket residue side chains are shown as sticks for comparison. **(M)** Sequence alignment of bridge helix and trigger loop N-terminus of eukaryotic RPB1, archaeal Rpo1N and bacterial RNA polymerase subunit β'. Hs, *Homo sapiens*; Dm, *Drosophila melanogaster*; At, *Arabidopsis thaliana*; Sp, *Schizosaccharomyces pombe*; Sc, *Saccharomyces cerevisiae*; Mj, *Methanocaldococcus jannaschii*; Bs, *Bacillus subtilis*; Ec, *Escherichia coli* (top) and H. sapiens POLR1A (Pol I), RPB1 (Pol II), and RPC1 (Pol III) (bottom). Threonine pocket residues are indicated with a red box. (**N**) Front view of sidetracked RNA base substituted *in-silico* for a guanosine (shown as red sticks). Red dashed line indicates steric clash between guanosine and RPB1 T857. Threonine pocket residues side chains shown as sticks

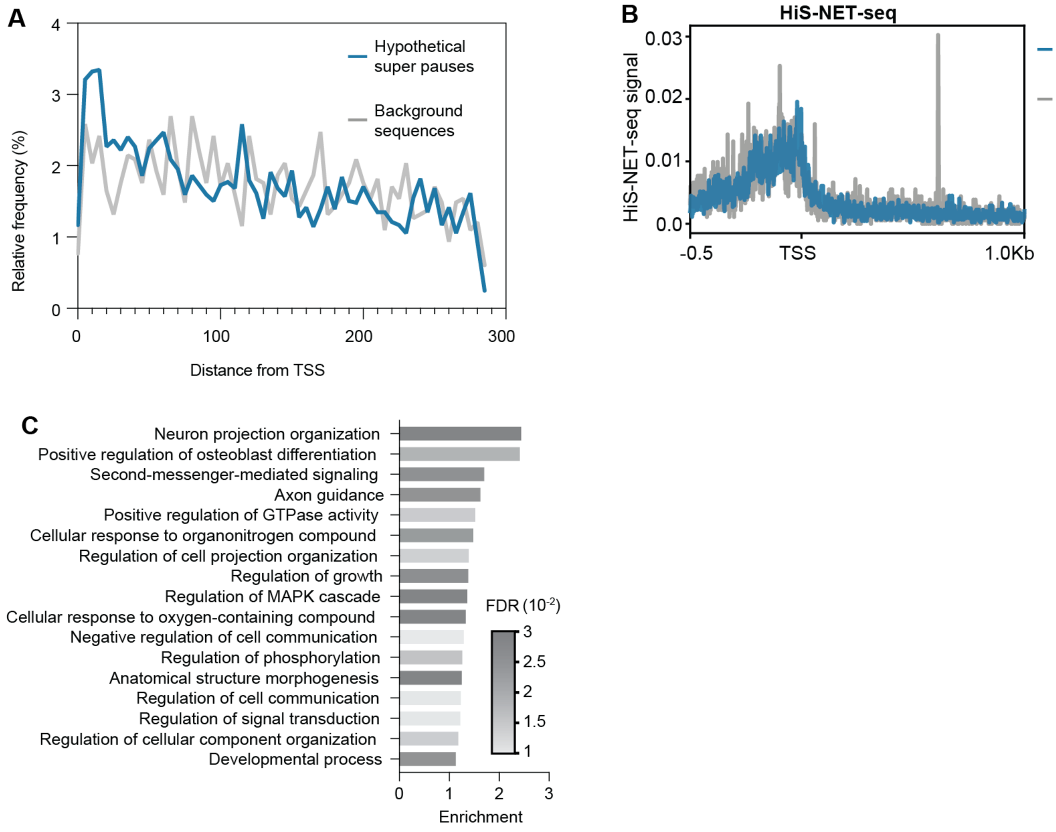


### Supplementary Fig. 10

**Supplementary Figure 10. Distribution and conservation of super pause sequences.**
(**A**) Distribution of hypothetical super pause sequences (n= 1273) (blue) and background sequences (gray) (n=1136) within the first 300 base pairs of 10,000 human protein coding genes. Values shown as percentage of the total number of detected sites. **(B)** Profile of genes with hypothetical super pause sites and background sites from HiS-NET-seq (**C**) GO term enrichment of genes with experimentally determined and hypothetical super pause sequences from panel A. FDR was used to correct for multiple hypothesis testing.
